# G-quadruplexes are promoter elements controlling nucleosome exclusion and RNA polymerase II pausing

**DOI:** 10.1101/2023.02.24.529838

**Authors:** Cyril Esnault, Encar Garcia-Oliver, Amal Zine El Aabidine, Marie-Cécile Robert, Talha Magat, Kevin Gawron, Eugénia Basyuk, Magda Karpinska, Alexia Pigeot, Anne Cucchiarini, Yu Luo, Daniele Verga, Raphael Mourad, Ovidiu Radulescu, Jean-Louis Mergny, Edouard Bertrand, Jean-Christophe Andrau

**Affiliations:** Institut de Génétique Moléculaire de Montpellier, University of Montpellier, CNRS-UMR 5535, 1919 Route de Mende, 34293 cedex 5, Montpellier, France; Institut de Génétique Humaine, CNRS-UMR9002, University of Montpellier, 141 rue de la Cardonille, 34396, Montpellier, France; Laboratoire de Microbiologie Fondamentale et Pathogénicité, CNRS-UMR 5234, Université de Bordeaux, Bordeaux, France; Laboratoire d’Optique et Biosciences, Ecole Polytechnique, CNRS, Inserm, Institut Polytechnique de Paris 91128 Palaiseau, France; Université Paris Saclay, CNRS UMR9187, INSERM U1196, Institut Curie, Orsay, France; LBCMCP, Centre de Biologie Intégrative (CBI), Université de Toulouse, CNRS, UPS, 31062 Toulouse, France; Laboratory of Pathogen Host Interactions, UMR CNRS 5235, University of Montpellier, Montpellier, France; Equipe labélisée Ligue Nationale Contre le Cancer, Montpellier

## Abstract

Despite their central role in transcription, it has been difficult to define universal sequences associated to eukaryotic promoters. Within chromatin context, recruitment of the transcriptional machinery requires opening of the promoter but how DNA elements could contribute to this process has remained elusive. Here, we show that G-quadruplex (G4) secondary structures are highly enriched mammalian core promoter elements. G4s are located at the deepest point of nucleosome exclusion at promoters and correlate with maximum promoter activity. We found that experimental G4s exclude nucleosomes both *in vivo* and *in vitro* and display a strong positioning potential. At model promoters, impairing G4s affected both transcriptional activity and chromatin opening. G4 destabilization also resulted in an inactive promoter state and affected transition to effective RNA production in live imaging experiments. Finally, G4 stabilization resulted in global reduction of proximal promoter pausing. Altogether, our data introduce G4s as *bona fide* promoter elements allowing nucleosome exclusion and facilitating pause release by the RNA Polymerase II.

## Introduction

Initially defined by analogy to bacterial transcription models (Jacob et al., 1964; Pribnow, 1975; Rosenberg and Court, 1979) and based on *in vitro* transcription assays from the pre-genomic era, eukaryotic core promoters were defined as *‘the minimal stretch of contiguous DNA sequence that is sufficient to direct accurate initiation of transcription by the RNA polymerase II machinery’* (Butler and Kadonaga, 2002). However and unlike in bacteria, eukaryotic promoters require nucleosome exclusion for the transcriptional machinery to be recruited. Beyond the concept of ‘core promoters’, eukaryotic ‘promoters’ can be characterized by wider sequence contexts that are highly divergent depending on the species. Nevertheless, these sequences carry in common to generate a well-positioned array of nucleosome (Jiang and Pugh, 2009; Radman-Livaja and Rando, 2010) and the ability to open chromatin. In mammals this later property is in part carried over by CpG islands (CGIs) (Deaton and Bird, 2011), characterized by large regions of high GC and CpG content that encompass a large fraction of promoters. Importantly, CGIs can open chromatin by default and in a transcription-independent manner (Fenouil et al., 2012).

Promoters are also highly enriched in potential DNA secondary structures, suggesting that these structures play a role in transcription regulation (Bansal et al., 2014). Among these, G-quadruplexes (G4s) are over-represented in regulatory regions. G4s are single-stranded and stable structures *in vitro* that consist in planar arrangement of guanines stabilized by K+ at physiological concentrations. They have been involved in number of nuclear processes and are present at over a million occurrence in the genome and more specifically at promoters. However, and to date it was unclear to what extent they could contribute as positive regulators of transcription since it was described that they either inhibit or repress transcription depending on their promoter context (Agarwal et al., 2014; Bochman et al., 2012; David et al., 2016; Smestad and Maher, 2015). Furthermore, while it was proposed that their formation *in vivo* could be dependent on high level of transcriptional activity (Hansel-Hertsch et al., 2016; Xia et al., 2018), the possibility that they could represent promoter elements on their own was never directly tested.

Here, we shed light on G4s as highly enriched mammalian core promoter elements. Using both predicted and experimental G4 data, *in vitro* and *in vivo*, we find them located at the deepest point of nucleosome exclusion at promoters correlating with maximum promoter activity in SURE assay. Furthermore, we found that G4s exclude nucleosomes both *in vivo* and *in vitro*, and display a strong nucleosome positioning potential. Impairing G4 formation by specific mutations at model promoters affected both transcription activity and chromatin opening. G4 destabilization also resulted in increased probability of promoters to be in an inactive state (OFF times) and affected transition to effective RNA production in live imaging experiments. Finally, G4 stabilization using ligands, resulted in global reduction of proximal promoter pausing by Pol II consistent with our live imaging observations. Altogether, our data introduce G4s as *bona fide* promoter elements allowing nucleosome exclusion and facilitating pause release by Pol II.

### G-quadruplexes are highly enriched at mammalian promoters and correlate with maximum promoter activity

Based on the initial knowledge that the TFIID general transcription factor (GTF) binds naked DNA *in vitro* in a window frame of 40-50 bp upstream and downstream of transcription start sites (TSSs) (Buratowski et al., 1989), *in silico* sequence analyses of core promoters were often restricted to this short window frame. However, these searches often yielded motifs poorly enriched, lowly conserved in evolution or highly degenerated (Haberle and Stark, 2018; Vo Ngoc et al., 2017). Because the most open areas of chromatin extend on average up to 100 bp, we performed a motif search on core promoters associated to open chromatin upstream and downstream (−100/+20) of experimental TSSs in three mammalian cell types (primary T cells, K562 and Raji cells - Figure 1A and S1; see also Table S1 for data sets used in this study). This analysis revealed a prominent motif consistent with SP1 binding site with additional G stretches in the 3 cell types. These stretches are highly compatible with the formation of G4s *in vitro*, using the G4Hunter (G4H) predictive algorithm (Bedrat et al., 2016) at various stringencies. To consolidate this result, we investigated the frequency of predicted G4s (pG4s) or other motifs identified and found overall that stringent G-quadruplex predictions (pG4s at G4H1.5 and 2.0 thresholds) (Bedrat et al., 2016) show very high frequency as well as a strong enrichment above control sequences (observed/expected) (Bagchi and Iyer, 2016; Fenouil et al., 2012). Enrichments of pG4s (20-45%) are also higher than those of Ets and NF-Y and far above the TATA box motifs (Figure 1A, S1 and Table S2). Their enrichment is similar to that of the BRE and SP1 motifs, both compatible with G4 formation, which show high overlaps at promoters (Figure S2A). Finally, we note that promoters containing pG4s tend to exhibit less of the other motifs (Figure S2B) suggesting that they could have more propensity to function autonomously.

**Figure 1:**
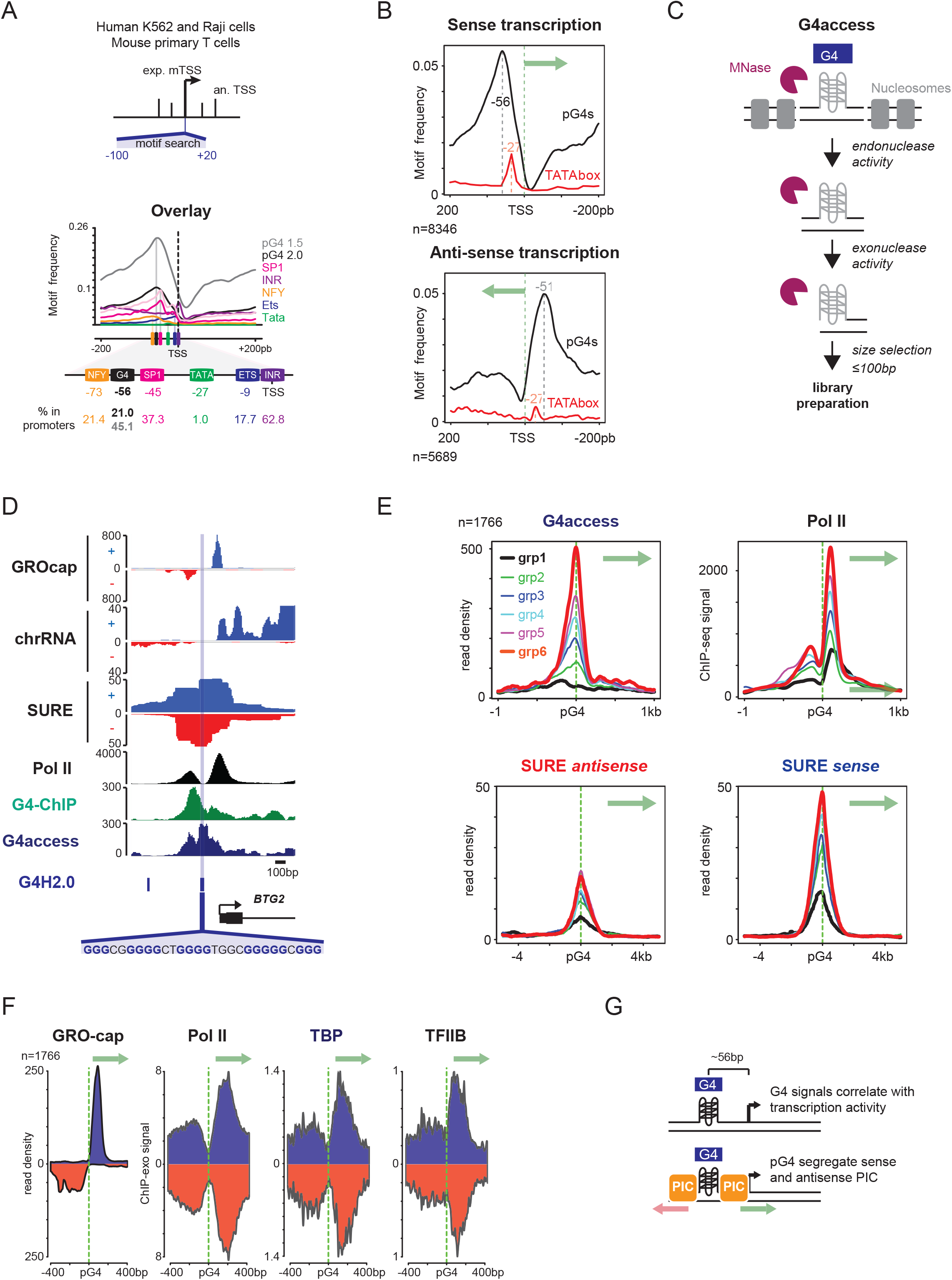
Predicted G-quadruplexes (pG4s) are highly enriched at promoters and correlate with maximum of mammalian promoter activity. (a) Promoter motif search around experimental TSSs highlights pG4 motifs in various cells. (Top) experimental strategy to determine promoter elements associated to experimental TSSs of expressed and annotated Refseq coding genes in our 3 model cell types (K562, Raji cells and mouse primary T cells); (bottom) motif distances to experimental TSSs, overlay of motif densities, relative representation at promoters and enrichment over control sequences are shown (INR= initiator). See also Figure S1 and Table S2 for detail of motif discovery analyses in 3 independent human and mouse cell types. (b) pG4s mark genomic areas comprised between sense and antisense promoters. Metaprofiles at promoters show enrichment of pG4s (G4H2.0) at 56 bp upstream of sense TSSs (n =8346) and at 51 bp of antisense TSSs for genes where divergent transcription initiation is detected (n=5689). See the heatmaps of Figure S1 for Pol II and short chromatin RNA profiles. A green arrow indicates the sense of transcription in each graph. (c) G4access principle. Chromatin is digested at moderate MNase level at which G4s show apparent resistance and are freed in subnucleosomal fractions. These are purified and subsequently subjected to library preparation and high-throughput sequencing. The details of the method are described in ref (Garcia-Oliver et al., 2022). (d) Example of a pG4 sequence fitting midpoint of upstream/downstream Pol II and nascent transcription (chrRNA and GROcap) at the *BTG2* promoter, experimental G4s (G4access and G4-ChIP), and maximum of promoter activity (SURE assay) in K562 cells are displayed. Sequence of the pG4 (G4H2.0) is indicated below the gene. (e) Experimental G4 signals (G4access) correlate with levels of transcription (Pol II ChIP) and SURE genome-wide promoter activity assay. Groups 1-6 correspond to increasing level of G4access signals. Green arrows represent the sense of gene transcription. Metaprofiles were centered on the G4 motif upstream of TSS as shown below the tracks. (f) pG4s are located at the midpoints of GROcap, Pol II, TBP and TFIIB GTFs using ChIP-exo datasets. See also Figure S3d-e for analyses of ChIP-seq in Raji and Mouse T cells. (g) Model of average pG4 locations as determined by G4Hunter algorithm upstream of TSSs and as a midpoint of upstream and downstream Pol II peaks. See Table S1 for data sets used, references and GEO accession numbers.

Next, we analysed the enrichment of pG4s upstream of sense and antisense transcription occurring at mammalian promoters. This showed that pG4s are found on average at positions at −56 and −51 of the TSS, respectively (Figure 1B). We note that while poorly enriched at promoters, when present, TATA boxes influence a far more directional and focused transcription (Figure S2C) as described previously (Bagchi and Iyer, 2016; Fenouil et al., 2012).

To further investigate whether G4s contribute to transcription initiation and promoter activity, we took advantage of four orthogonal approaches for G4 formation assessment, including G4access a technology we recently developed in our laboratory (Garcia-Oliver et al., 2022). G4access is an antibody-independent technology alternative and complementary to G4 ChIP. In brief, this method is based on moderate nuclease digestion of chromatinized DNA and allows G4 formation assessment genome-wide in the chromatin context (Figure 1C). We also used G4 ChIP (Hansel-Hertsch et al., 2016; Mao et al., 2018), mapping G4s in living cells, G4seq (Chambers et al., 2015) globally assessing G4s *in vitro* and ss-DNA-seq (Kouzine et al., 2013) mapping single stranded DNA genome-wide. We performed G4access in Raji and K562 cells and processed published data sets for the other methods whenever available (Figure S2D-E). In this analysis, we ranked the promoters containing predicted G4s (G4Hunter>2.0) by increasing experimental G4 signal. G4seq and ssDNA-seq in Raji cells were used to further validate the selection of promoters with folded G4s. G4access and G4-ChIP outputs of G4 measurement are very comparable (Figure S2D-E). ssDNA-seq globally confirmed G4 formation in Raji cells while G4-seq, a technique that maps G4s formed *in vitro* on naked genomic DNA (Chambers et al., 2015), validated that selected sequences can form G-quadruplexes *in vitro*. Since G4-seq monitors genomic G4 formation outside of nuclear context, signals of the 6 groups originally defined from low to high G4 formation in living cells remain largely unchanged. This further indicates that G4access and ChIP do map G4s in the context of chromatin in living samples.

We then compared G4 formation to data sets (Table S1) monitoring nascent transcriptome and large-scale measurement of promoter activity assay (SURE) (van Arensbergen et al., 2017) (Figure 1D-E). As illustrated for the BTG2 promoter in K562 cells, both predicted and experimental G4 signals correspond to the midpoint of promoter divergent transcription and maximum promoter activity by SURE (Figure 1D). To establish this statement globally, we analysed the correlation of G4access at pG4 promoter locations with that of SURE and Pol II (ENCODE) and found that both increase with experimental G4 levels (Figure 1E). In addition, we observed that pG4 locations overlapped with the midpoints between sense and anti-sense transcription initiation mapped by ChIP-exo or ChIP-seq of Pol II, TBP, TFIIB (Pugh and Venters, 2016) and by GRO-CAP (Core et al., 2014) (Figure 1F and S2G). Finally, we also observed that promoter pG4s localise upstream of R-loops (Figure S2F) which form where Pol II and GRO-cap signal start to rise, on the side of the pG4s (Figure 1F).

Collectively, these analyses show that pG4s represent major motifs of extended core promoters, located on average at a relatively fixed position from TSSs. Moreover, G4s correlate with both promoter activity and midpoints of divergent transcription (Figure 1G).

### G4 forming sequences carry an intrinsic ability for nucleosome exclusion

Apparent Nucleosome Depleted Regions (NDRs) are hallmarks of core promoters in eukaryotic cells allowing space for PIC recruitment (Andersson and Sandelin, 2020; Haberle and Stark, 2018). To understand the link between G4s/pG4s and nucleosome positioning, we performed nucleosome mapping by MNase-seq in Raji cells and re-analysed published datasets, for nucleosomes and active epigenetic marks in our two other mammalian models (Table S1). Strikingly, we found the center of NDRs overlapping with pG4s at a very large fraction of promoters, including at the BTG2 promoter described above (Figure 2A). We then investigated all active promoters that contain pG4s and confirmed that pG4s are found at the deepest points of NDRs globally (Figure 2B, Figure S2H). By comparing increasing G4 signals to promoter opening, we observed less opening in the absence of G4access signal (group1). Conversely, deeper and larger NDRs were present when G4access was present (group 2-5), suggesting a threshold effect in the G4access signal. Active histone marks increased together with G4 strengh (group 1-6), indicating overall chromatin opening and modifications depending on the G4 formation (Figure 2B). At inactive promoters, the presence of predicted G4s also hallmarked Polycomb-deposited H3K27me3 inactive chromatin mark (Figure S2I). This set of promoters carry the hallmark of CpG islands, with strong GC and CpG content as expected for Polycomb signal. Since SP1 motifs are also highly enriched at promoters (Reed et al., 2008) and represent half of a pG4 motif (Huppert et al., 2008), we questioned whether SP1 binding or motif could influence the observed chromatin opening by G4s. We analysed pG4 promoters with or without SP1 binding and observed similar NDR formation, reasonably allowing to exclude that SP1 would be responsible for the pG4 property (Figure S3A). Similarly, promoters that did not carry the canonical or non-canonical SP1 motifs did also show openness at G4 motifs (Figure S3B-C). In this case, we observed residual SP1 binding indicating that SP1 might directly bind G4s in accordance with published observations (Lago et al., 2021; Raiber et al., 2012).

**Figure 2:**
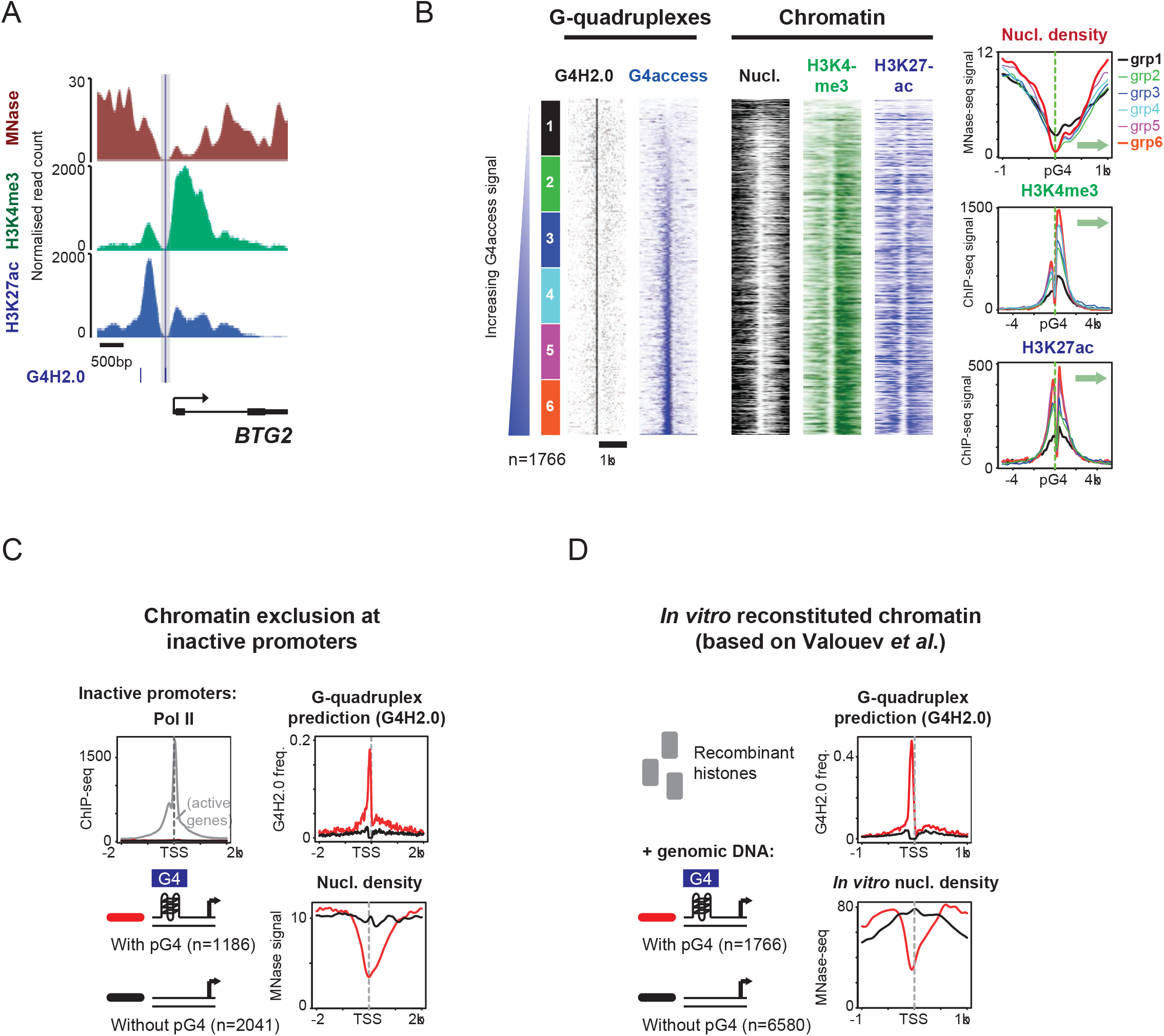
G4s promote nucleosome exclusion at active and inactive promoters, *in vivo* and *in vitro* (see also Figure S2 and S4). (a) Example of a pG4 sequence fitting the maximum of the nucleosome depleted regions (NDRs) at the Btg2 promoter in K562 cells using MNase-seq, and ChIP-seq of active chromatin marks (H3K4me3, H3K27ac). (b) Experimental G4 signals correlate with levels of nucleosome depletion and with active histone modification marks. Promoters that harbour strong G4 predictions using G4Hunter algorithm G4H2.0 were ranked by G4access signal as depicted in the heatmap, corresponding heatmaps of G4-ChIP and G4H2.0 predictions are also shown (left). Promoters were split in 6 groups, as in Figure 1E. Metaprofiles of nucleosome densities (MNase-seq) and of H3K4me3 and H3K27ac of all groups are displayed (right). See also Figure S2H for other cell types. Green arrows represent the sense of gene transcription. (c) pG4 promote nucleosome exclusion at inactive promoters in K562 cells (ENCODE). Transcriptionally inactive promoters were split in groups with or without G4H2.0 prediction. See also Figure 3 for pG4 influence on nucleosome positioning at inactive intergenic regions and Figure S4A after transcription inhibition with a-amanitin. (d) pG4-containing promoters have intrinsic nucleosome exclusion properties on *in vitro* reconstituted chromatin (analysed from (Valouev et al., 2011)). The promoter selections are based on K562 active promoters shown in Figure 1. See also Figure S4B-C for pG4s effect on nucleosome exclusion properties on *in vitro* at inter and intragenic regions.

Next, we tested whether nucleosome exclusion at pG4s was dependent on transcriptional activity. For this, we analysed nucleosome densities at inactive promoters (see methods) that were separated into 2 groups, with or without pG4s. Interestingly, only pG4-containing promoters showed significant nucleosome exclusion (Figure 2C). We further confirmed that transcription is not required for nucleosome exclusion at pG4s of active promoters by inhibiting Pol II transcription with a-amanitin without substantial loss of NDRs (Figure S4A). To further infer and demonstrate the direct link between pG4 sequences and chromatin opening, we made use of reconstituted nucleosomes *in vitro* (Valouev et al., 2011). In this assay, chromatin was reconstituted using human genomic DNA and recombinant histone before MNase digestion. As for the *in vivo* analyses, promoters were split in 2 groups, with or without strong G4 predictions (Figure 2D). This analysis revealed that only the pG4-containing group associates with nucleosome exclusion. Thus, pG4 DNA carry the intrinsic ability to exclude nucleosomes since *in vitro*, in absence of any other transcription factor or proteins, we could observe this property. It also indicates that nucleosome occupancy and G4 formation are mutually exclusive. This ability to exclude nucleosome is similarly observed for the SP1 unbound promoters (Figure S3). Together, our data show that at both active and inactive promoters, G-quadruplexes promote intrinsic nucleosome exclusion *in vivo* and *in vitro*.

### G4 forming sequences are nucleosome organizers

Next, we investigated chromatin opening and organisation around G4 predictions at non-promoter locations, including intergenic regions. These locations are mainly not transcribed and yet, pG4s are associated to nucleosome exclusion (Figure 3A) indicating that these sequences carry this property over the whole genome and not only at promoters. Further investigation of *in vitro* reconstituted chromatin (Valouev et al., 2011) at all genomic locations including promoters, intergenic and intragenic regions demonstrated their intrinsic abilities to evict nucleosome globally (Figure S4B-C). Nucleosome exclusion was found at over 90% for promoter pG4s, and over 80% for intra and intergenic pG4s (Figure S4C). Interestingly, the regions showing the largest nucleosome exclusion areas also showed the highest G4 prediction densities (upper part of the heatmaps), possibly because their presence in clusters increase chances of G4 formation. These observations are consistent with the high stability of G4 single-stranded DNA structures (Guedin et al., 2010; Sen and Gilbert, 1988) which makes them incompatible with nucleosome formation.

**Figure 3:**
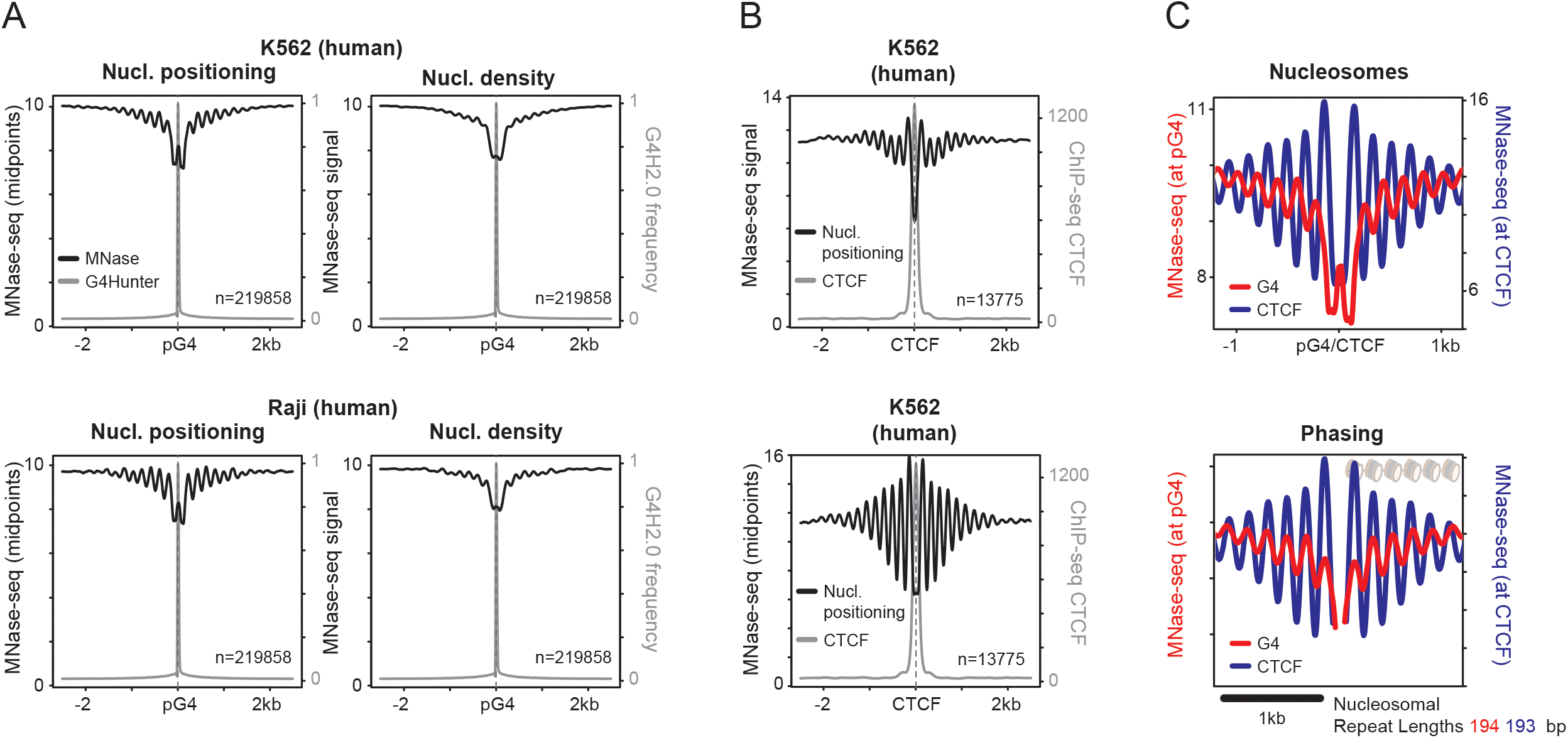
pG4s organise nucleosome at intergenic regions (IGRs). (a) pG4s mark the center of organised nucleosome arrays at IGRs in K562 (ENCODE) and Raji (this study). All G4H2.0 predictions from IGRs are shown. Heatmaps of nucleosome organisation mapped by MNase-seq is displayed. Nucleosome positioning (see methods, left), MNase-seq density (right). (b) Nucleosomal organisation at CTCF sites in K562 cells. Metaprofiles of MNase-seq signal (Top), positioning (Bottom) and CTCF ChIP-seq (GSE30263) at the 13775 identified binding sites in K562 cells. (c) Nucleosomal organisation at CTCF and pG4 show similar nucleosome phasing at IGRs. (Top) Overlay of metaprofiles of MNase-seq at IGRs centred either on pG4s or CTCF ChIP-seq sites in K562 cells; (Bottom) the two nucleosomes surrounding CTCF or pG4 sites (−1/+1) were aligned to compare similarity of nucleosome phasing (observed nucleosome repeat lengths of 193 and 194 nt, respectively).

To infer how nucleosome behave around G4 forming sequences, we analyzed both nucleosome densities and midpoints around intergenic G4 predictions in our model cell lines. Midpoints analyses allow to better assess if given sequence locations display positioning properties (Figure 3A). This clearly revealed a high level of positioning associated to pG4s in the model cells. The periodicity of observed nucleosome positioning is highly similar to nucleosomal organization around specific pioneer transcription factors (Barozzi et al., 2014) or the insulator factor CTCF(Fu et al., 2008) showing an almost identical nucleosomal repeat length (Figure 3B-C), in line with previous observations (Kouzine et al., 2017). Hence, our results highlight the association of pG4s with open chromatin regions at promoters and at IGRs where pG4s also associate with nucleosome array organization. Although experimental G4 signals correlate with transcription, pG4/G4 driven nucleosome depletion appears independent of transcription. This observation is consistent with results described recently by us and others (Garcia-Oliver et al., 2022; Shen et al., 2021). Our results support the role of a novel and previously unappreciated role for G4s as global chromatin organisers at transcribed and un-transcribed regions.

### Increased chromatin opening at CpG islands containing pG4s

Since CpG islands (CGIs) are able to promote nucleosome depletion at promoters (Fenouil et al., 2012), we wondered what was the contribution of pG4s in this process. To address this question, we considered all human CGI annotations containing or not strong G4 predictions (using G4H2.0, Figure 4). We analysed genomic features associated to experimental G4s: nucleosome positioning, active chromatin marks, transcription and promoter activity. As expected, G4access and ChIP-seq show stronger signals at CGIs harboring pG4s. In addition, active histone marks (H3K4me3 and H3K27ac) and Pol II are also enhanced in this class (Figure 4A-B). Furthermore, nucleosome occupancy exhibits wider and deeper chromatin opening at pG4-containing CGIs. Finally, the analysis of promoter activity by SURE assay confirmed that G4-containing CGIs have a ~1.5 time higher promoter activity. To further validate our results, we confirmed our analyses on a set of CGIs of same lengths and CG contents (Figure S5A) with a more stringent selection of non-G4 forming sequences. The G4 forming sequences were considered with a G4Hunter score> 1.5 and non-forming sequences for a G4Hunter score < 1.2. This analysis confirmed the association of G4 forming sequences to more open chromatin and active transcription and epigenetic marking within CpG islands (Figure S5B-C). Taken together, our observations support the notion that features characteristic of promoters are enhanced in the presence of both experimental and predicted G4s at CGIs and that G4s might represent essential determinants of CGI’s ability to exclude nucleosome.

**Figure 4:**
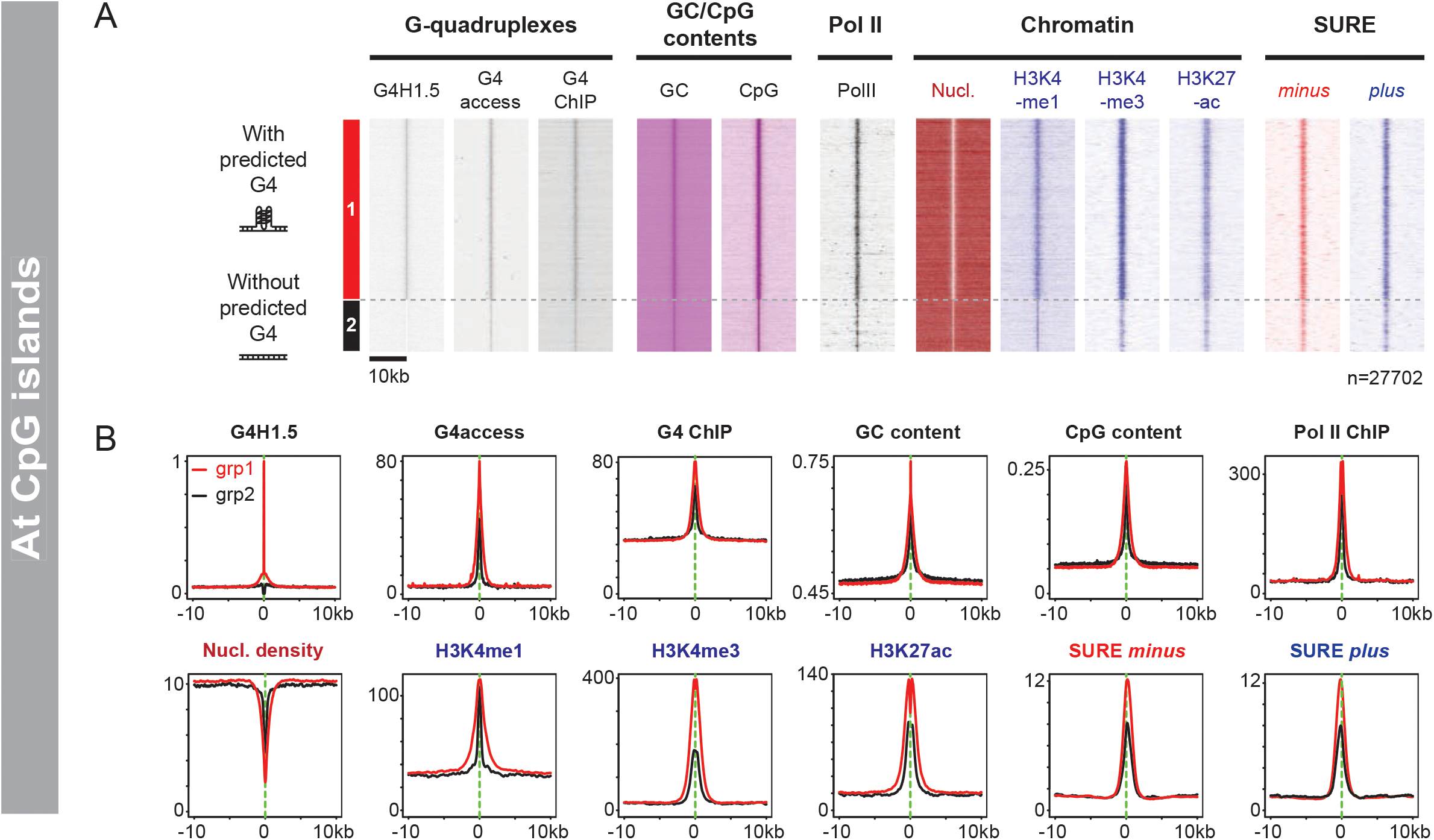
pG4s and experimental G4s hallmark nucleosome exclusion at CpG islands (CGIs) (see also Figure S5). (a) CGIs with pG4s have enhanced chromatin opening, histone modifications and transcription activity. Heatmaps of the 27702 human CGIs were split in two groups with (n=21520) or without (n=6182) pG4 (G4H1.5) annotations. Corresponding signals for pG4, G4access, G4 ChIP, GC and CpG contents, Pol II, nucleosomes (MNase-seq), chromatin marks and SURE promoter activity are shown as indicated. (b) CGIs with pG4s have deeper chromatin opening, increased active histone modifications and promoter activity. Metaprofiles of all marks at CGIs with or without pG4 displayed in the heatmaps from (a). Further selection and controls for this anlaysis are presented in Figure S5.

### G4 mutations at promoters result in decreased transcription in single cells

To fully demonstrate that G4s are promoter element that extend the concept of the core promoter, we mutated G4s in model promoters. We inserted G4-containing mouse promoters in the human genome in Hela cells using a Flip-in system as described previously (Tantale et al., 2016a). The promoters were located upstream of a reporter gene containing 256 MS2 repeats and allowing single cell measurement of transcription by smRNA FISH (Figure 5A). We chose promoters that contained a strong G4 prediction that was also verified experimentally using G4access (Garcia-Oliver et al., 2022) or G4 cut&Tag (Lyu et al., 2022) in mouse ES cells (Figure S6A-E). For 3 out of 5 models (Taok1, Pkm and Klf6), the sequences considered did not contain any TATA box, and for 3 of them no SP1/GC box site (Taok1, Pkm and Klf6). We designed G4 mutations that minimally affected the promoter primary sequence while impairing the G4 potential (Figure 5B). We also verified that the structure was abolished *in vitro* using 3 independent biophysical assays that included circular dichroism, TDS and IDS (thermal and isothermal differential spectra, Figure 5B). In the case of the Pol2ra, we performed 2 independent mutations that affected differentially the G4 potential assessed by the G4hunter algorithm (Figure 5B and table S3), one of which respecting the GC content of the sequence. To quantify the transcriptional output of the model promoters, we performed single cell measurements using smRNA FISH over hundreds of cells. The data presented in Figure 5C shows G4 mutant’s transcription as compared to their WT promoter counterpart. The level of nascent RNA reduction ranked from 30 to 80% in the mutants, indicating that the G4 mutations had a significant effect on transcription of the core promoter. Interestingly also, inverting the G4 orientation in the promoter did not impact on its activity in the case of the Eef1a1 model (Figure S6F), suggesting that the G4 functions as promoter element in an orientation independent manner.

**Figure 5:**
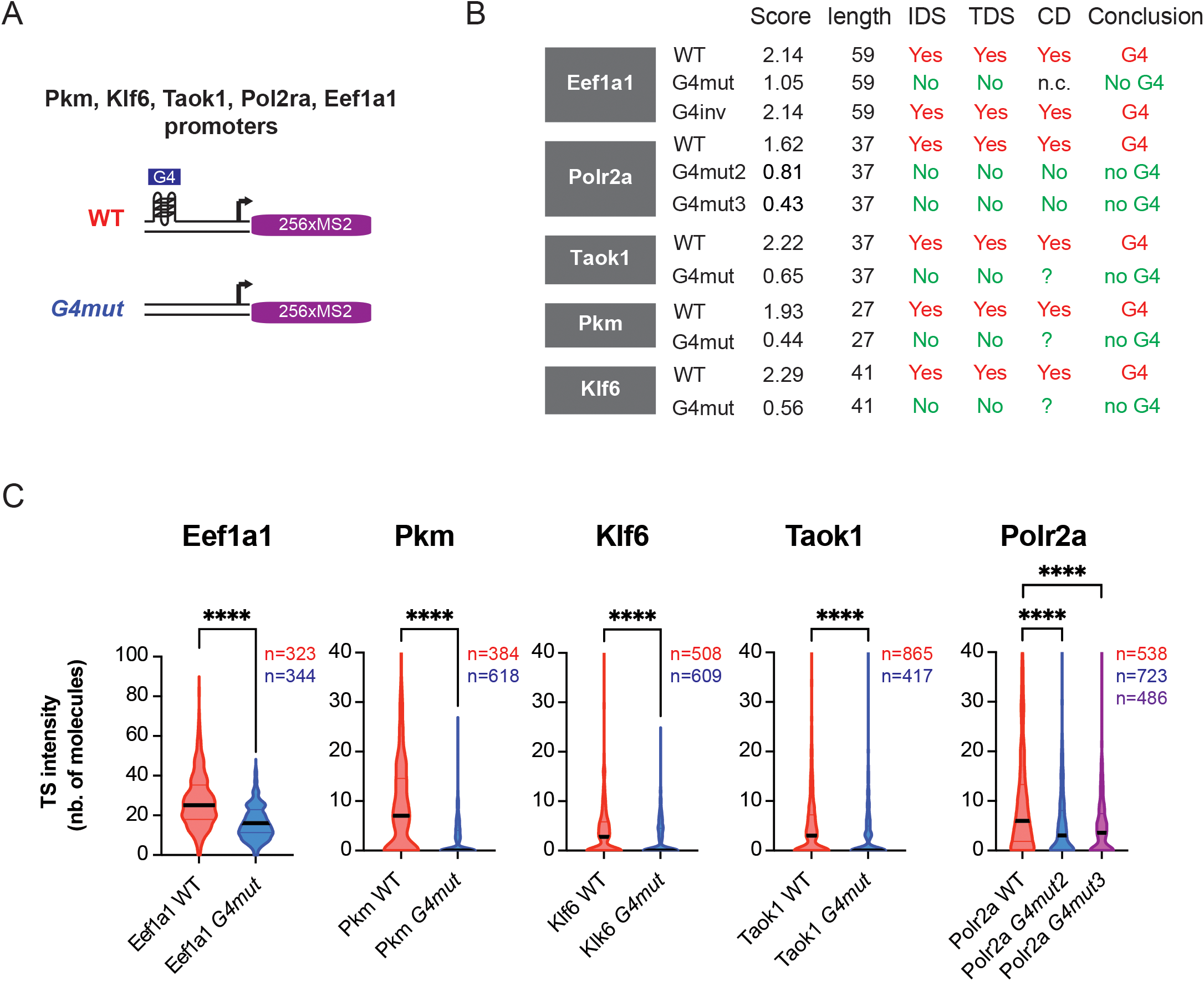
G4 mutations impair promoter activity in single cells (see also Figure S6 A-F and Table S3). (a) Scheme of integrated reporter constructs for the indicated mouse G4-containing promoters used for smRNA FISH. These constructs were integrated in human genome (Hela) using a Flp-In strategy. Experimental G4 signals for the mouse promoters in ES cells are shown in Figure S6A-E. (b) Sequence features and G4 structural assessment in the model promoters indicated in (a). The G4 scores determined by the G4Hunter algorithm, reflecting stability and likelihood of formation are indicated for WT and mutant sequences (see Table S3 for the individual sequences). Three independent assays were performed to conclude for G4 formation *in vitro* on the oligonucleotide (last column). (c) Quantification of smFISH images of MS2 reporter activity of Ee1fa1, Pkm, Klk6, Taok1 and Polr2a WT and mutant promoters. Mann-Whitney tests were used (ns P > 0.05; **** P ≤ 0.0001). For Polr2a the two mutants have moderate (mut2) or strong mutations (mut3) (see panel b). Sequences of all promoters are provided in Table S3.

Together, our data show that G4 mutations impair transcription quantitatively and establish that G4-forming structures most likely function as promoter elements in an orientation independent manner.

### Modelling transcription in presence or absence of G4 and TATA box elements

To decode the direct effect of G4s as promoter elements on transcription, as compared to that of the TATAbox, we performed further mutational analyses in the context of the Eef1a1 promoter that contains both a canonical TATA box and a stable G4. We mutated the G4, the TATA, or both elements (Figure 6A). Minimal substitution of the TATA box and G4s were used to affect the functions of the elements. We first analysed their ability to promote transcription, to recruit the pre-initiation complex and to assemble chromatin. Our smRNA FISH of MS2 reporter results show that both G4 and TATA elements are required for full activity (Figure 6B). We found that pG4 mutation display more pronounced effects (~60%) while less impact of the TATAbox was observed and an additive or synergic effect seems at play in the double mutant.

**Figure 6:**
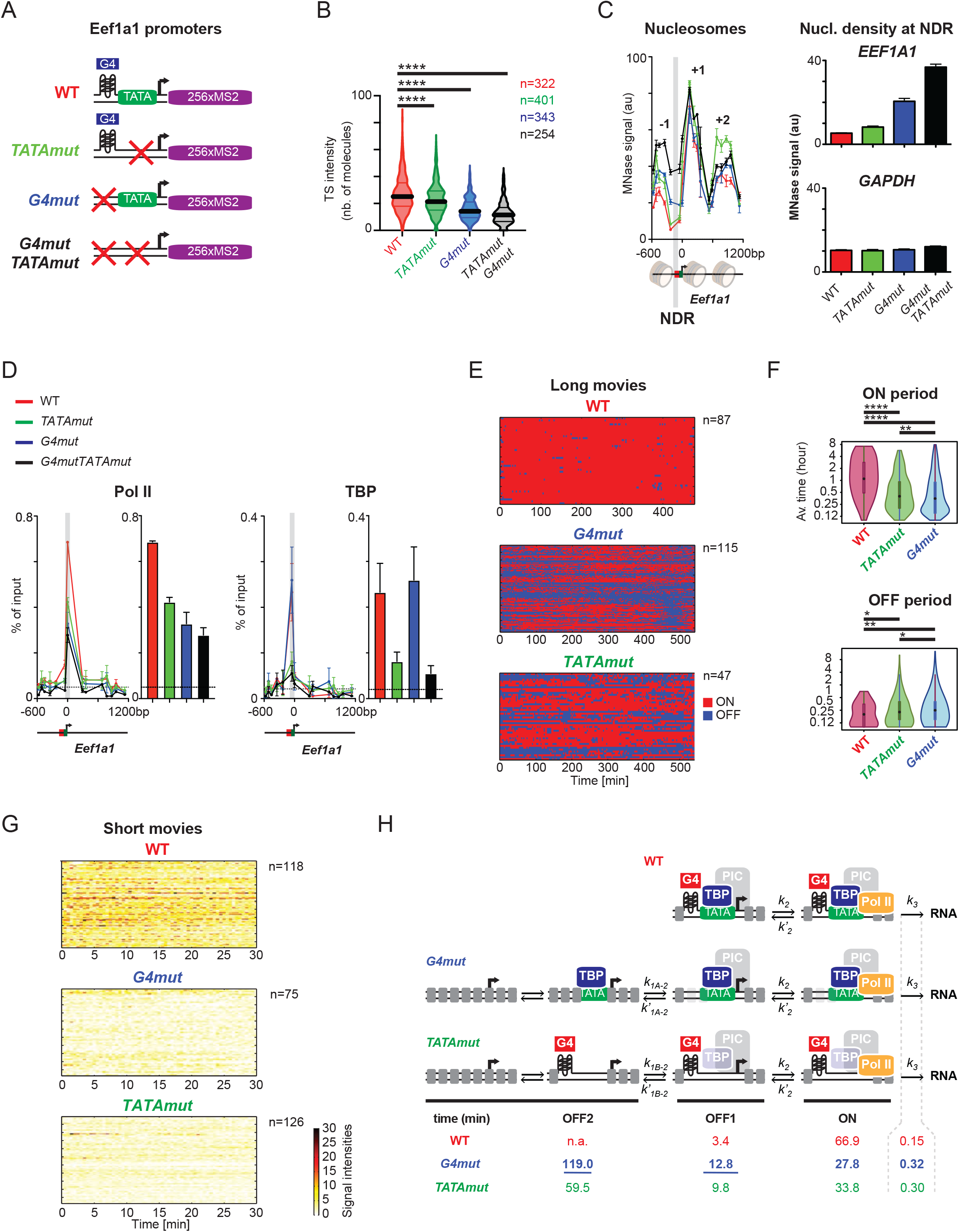
G4 mutations at Eef1a1 model promoters decrease transcription and increase nucleosome density at NDRs, while increasing promoter OFF times in single cells (see also Figure S6 G-I). (a) Scheme of Eef1a1 mouse model promoters inserted in human HeLa cells with indicated mutations (see also Table S3 for the sequences) (see also Table S3 for the sequences). (b) G4 forming sequences and the TATAbox regulate gene expression from the Eef1a1 model promoter in single cells. Quantification of smFISH images of MS2 reporter activity of WT and mutant Eef1a1 promoters. Representative smFISH images are shown in Figure S6G. Mann-Whitney tests were used. (c) Eef1a1 G4 mutations result in increased nucleosome density at the apparent nucleosome depleted region (NDR) location. MNase signal of WT and mutant promoters was quantified by qPCR (n=3; means ±s.e.m), the NDR location is highlighted in grey. Quantifications of the nucleosome density signal variations at NDR are shown on the right of the graph, together with a GAPDH control. (qPCR oligonucleotides used in are presented Table S4) (d) Pol II and TBP recruitment at Eef1a1 model promoter and mutants. ChIP were assayed and quantified by qPCR (n=3; means ±s.e.m). Pol II is affected in all mutant contexts while TBP is impaired specifically in the TATAmut. (qPCR oligonucleotides used in are presented Table S4) (e) Heatmaps of permissive (red) and non-permissive (blue) transcription periods for WT and mutant Eef1a1 promoters in single cell live imaging (Long movies, 8h-9h, with stacks every 3 min). Each line represents an individual cell assessment. Representative movie images and promoter threshold are shown in Figure 6G. (f) Violin plots of ON and OFF period average duration in WT or mutated Eef1a1 promoters measured in long movies). Computed Pvalues (Mann-Withney test) are as follows: ****<1e^-4^, ***<1e^-3^, **<1e^-2^, *<5e^-2^ (g) Heatmaps of the density of polymerases (number of polymerases every 30s) for WT and mutant Eef1a1 promoters in single cell live imaging (short movies, 30 min, with stacks every 30s). Each line represents an individual cell assessment. (h) Interpretative scheme of the signal deconvolution and time constant analysis derived from the long and short movies. While WT transcription can be described in 2 main steps both TATA and G4 mutants require at least 3 steps. The G4 mutant has limiting OFF2 and ON state (see also Figure S6H-I and methods for mathematical modelling).

To infer how the G4 and TATAbox influence promoter activity and transcription, we then analysed nucleosome organization and PIC assembly at these promoters in bulk assays. MNase assay coupled to PCR analyses revealed that mutating the G4 sequence led to increased nucleosome density at the level of the NDR location, that also corresponds to the G4 structure coordinate (Figure 6C). Remarkably, this effect was not observed in the TATA box mutant. The double mutant showed synergic effects of the elements, consistent with nascent transcription data. These results suggest that the pG4 is the main element controlling Eef1a1 promoter opening and that the TATAbox influence could only be seen in the TATAmutG4mut context. We next monitored PIC recruitment using Pol II and TBP ChIP qPCR assays. Interestingly, while Pol II recruitment was reduced in all mutants, TBP binding was only impaired when the TATAbox was mutated but not in the G4mut (Figure 6D). We conclude that the primary effect of the pG4 mutation is to restrict chromatin and Pol II accessibility but not TBP recruitment, while the TATAbox mutation affects TBP and ultimately Pol II recruitment (Figure 6A-D).

Next, we analysed the Eef1a1 promoter dynamics. We examined transcription in living single cell by imaging an MCP-GFP fusion protein. We recorded long movies on the WT, TATAmut and G4mut Eef1a1 cell lines, by taking 3D image stacks every 3 min during 8 h (see representative examples in Figure S6G). Since a single polymerase remains more than 3 min at the reporter transcription sites (Tantale et al., 2016a), and since the sensitivity of this assay allows single polymerase tracking, every single transcription event can be detected. We quantified the brightness of transcription sites in hundreds of cells (see methods) and quantified permissive (ON) periods, from which Pol II regularly initiate transcription, from inactive (OFF) periods (Figure 6E). These analyses clearly show that both TATA and G4 mutants displayed shorter ON and longer OFF periods. Moreover, OFF periods were slightly longer for the G4 mutant as compared to TATAmut (Figure 6E-F). To complete this picture, we also recorded short movies at high temporal resolution (one stack every 3 seconds), to model the entire dynamics of the Eef1a1 promoter (Figure 6G). By performing mathematical modelling of the distribution of initiation events and interpolating the signal intensities (Figure S6H) (Tantale et al., 2021), we were able to show that the WT promoter is described by a two states promoter model (ON and OFF). In contrast, both G4 and TATA mutants require three states model to describe the experimental data, with the additional promoter state being a long-lived inactive state (~2h lifetime; Figure S6H-I). Given the results of the biochemical analysis of the mutant promoters, the additional state likely corresponds to a TBP unbound state for the TATAmut, and a closed chromatin state for G4mut. Moreover, examination of transcription initiation rates derived from the models (k3), indicates a slower transition into processive elongation for the G4 mutant (0.3 vs 0.15 seconds). These live cell kinetic data are consistent with the idea that the G4 mutations severely limit chromatin opening. In addition, changes in the k3 constant (Figure 6G, S6I) suggest that G4 mutations also slow down promoter escape and/or Pol II pause release (Figure 6H).

Our observations on the model Eef1a1 promoter (summarized in Figure 6H) lead us to propose that the role of G4s at promoters is to promote and maintain nucleosome exclusion as a prerequisite for stable PIC recruitment. In contrast, when the TATAbox is mutated, formation of an active promoter state is also impacted due to defects in TBP and PIC recruitment. Collectively, our reporter experiments in bulk and single cells further support the role of G4s/pG4s as promoter elements, conditioning nucleosome exclusion and the rates of promoter transition toward an active state competent for transcription.

### G4 stabilization results in Pol II promoter proximal pause release

To investigate the influence of G4 stabilization on transcription at the global, we treated human Raji cells with Pyridostatin (PDS), a well-known G4 ligand (Rodriguez et al., 2008). This ligand stabilizes G4’s structure by limiting the transition from G4-structured to unstructured ssDNA or ds B-DNA (Rodriguez et al., 2008). Therefore, and since promoters are highly enriched in pG4s, their stabilization could either positively or negatively impact transcription. To address this question, we monitored PDS effects on Pol II densities and nascent transcripts, using ChIP-seq and chrRNA-seq, respectively. We used short time points of treatment (10 to 60 min) to avoid indirect effects (Olivieri et al., 2020; Rodriguez et al., 2008) resulting from the appearance of double strand breaks at later time points (Figure S7A). In these assays, we observed that PDS led to changes in Pol II profiles after only 10 minutes of treatment, with increased signal in gene bodies while signal at promoters was decreased. Examples in Figure 7A illustrate this effect for model genes in which Pol II release is illustrated by either promoter density decrease, gene body increase or both (the sequence of the G4 upstream of TSS is indicated). These effects most likely reflect a general promoter proximal pause release of Pol II. We confirmed a global decrease of Pol II pausing by computing apparent pausing scores at the genome-wide level (Figure 7B and Figure S7B). The effect was found more pronounced for a subset of 556 genes (see methods). Interestingly these genes tend to display slightly less stable G4s suggesting that the ligand preferentially act on weaker structure (Figure 7C) in agreement with our recent observation using the G4access procedure (Garcia-Oliver et al., 2022). This selection also appeared as enriched in mRNA splicing functions (Figure S7C). Consistent with pause release, Pol II average profiles at this subset show a decrease around TSSs and an increase of Pol II density over gene bodies (Figure 7D-E). This was further confirmed by nascent chrRNA-seq analysis (Figure 7F-G). We also analysed later time points, showing that, although reduced pausing is still visible at 30 min, the impact of PDS is partially reversed after 60 min over gene bodies (Figure S7B, middle panel). Altogether, our observations suggest that G4 stabilization results in increased ability for Pol II to escape from pause states. This is also consistent with our mathematical modelling of the eef1a1 model promoter where G4 mutations affected efficiency of Pol II released from the promoter.

**Figure 7:**
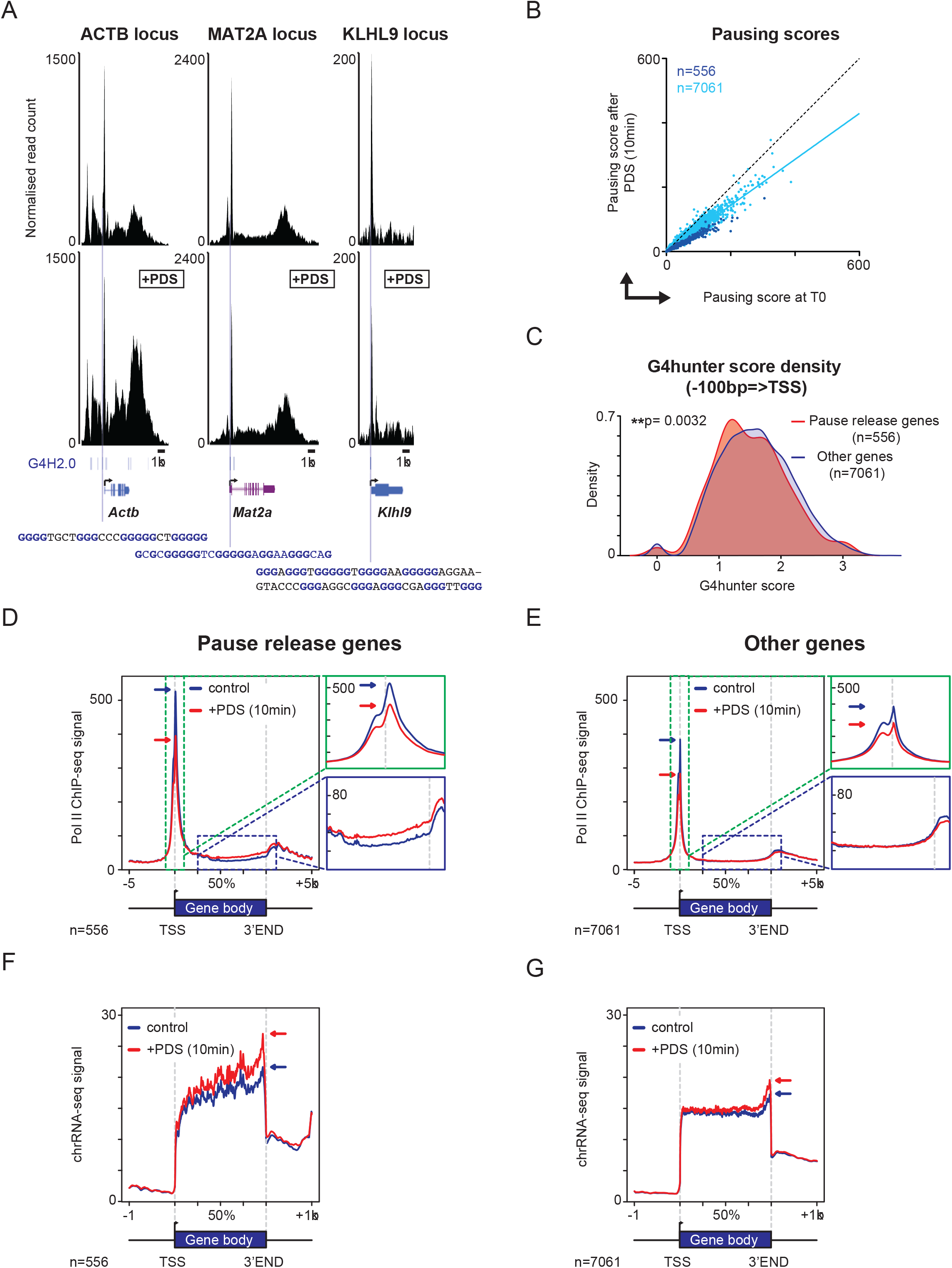
Stabilization of G4 by ligand results in global pause release by Pol II. (see also figure S7) (a) Pyridostatin (PDS) treatment (10 min) results in Pol II release from the promoter area and reduction of the +1 nucleosome at the ACTB, MAT2a and KHLH9 loci. Tracks of Pol II ChIP-seq and MNase-seq (zoomed around TSS) before and after PDS treatment are shown. G4 predictions and the sequence of the pG4 upstream of the TSSs associated with the NDRs are indicated and are shown below the tracks (G4H2.0). (b) Pausing scores are globally reduced at the genome-scale following PDS treatment. Scatter plots comparing pausing scores before and after PDS treatment (10 min) is shown. Genes with enhanced Pol II signal at gene bodies (Pvalue <0.05) are highlighted in dark blue (n=556). The light blue slope represents the linear regression curve of the 7061 other points. The whole kinetic analysis is presented in Figure S7B. (c) G4Hunter scores distribution of pause release genes and others. (d) Metaprofiles of Pol II ChIP-seq signal at selected genes with decreased pausing scores (n=556) depict reduced promoter and increased gene body occupancies. (e) Metaprofiles of Pol II ChIP-seq signal at control genes following PDS treatment. (n=7061). (f) Metaprofiles nascent ChrRNA-seq genes at pause release genes. (n=556). (g) Metaprofiles nascent ChrRNA-seq genes at other genes. (n=7061).

## Discussion

Our study has shown that G4 forming sequences are highly enriched in extended mammalian core promoters. We propose a novel function for these elements that is to intrinsically exclude nucleosomes, possibly defining one essential property of promoters *in vivo*. The mechanism of nucleosome exclusion by G4s could be simply explained by the incompatibility of stable single-strand DNA (ssDNA) formation and its incorporation into stable nucleosomes. Since G4s are not significantly present in all eukaryotic promoters (Marsico et al., 2019), other secondary structures or sequence context could have the same role in other organisms, for example AT stretches in yeast (Kaplan et al., 2010; Segal and Widom, 2009). This G4 property is however in contrast to previous observation proposing that it is transcription that stimulates G4 formation based on *in vitro* transcription (Xia et al., 2018) assays or transcription activation *in vivo* (Hansel-Hertsch et al., 2016) but in line with more recent observations (Shen et al., 2021), including work from our laboratory (Garcia-Oliver et al., 2022). In this work (Garcia-Oliver et al., 2022), we show that transcription inhibition results in maintenance, but reduction of G4 signal indicating that transcription does not precede G4 formation at promoter but that it does increase or stabilize its structure. Taken altogether, our data plead for a model in which G4s are formed prior from/or in the absence of transcription since pG4s/G4s are detected at NDRs *in vitro* or at silent promoters and since transcription inhibition does not significantly alter NDRs at pG4 locations. The presence of frequent transcription factor binding site (TFBS) motifs in addition to pG4 elements also opens the possibility that PIC recruitment *in vivo* does not directly rely on motifs that recruit the GTFs but rather on various already bound TFs, possibly in combination to co-activator, to allow further recruitment of the PIC. Arguing also for this model is the observation that TFIID does not contact DNA on its own upstream of TSSs on TATA-less promoters (Burke and Kadonaga, 1996; Parry et al., 2010), which represent the vast majority of promoters. Furthermore, TFIID recruitment was previously reported to occur through direct chromatin contacts (Bhuiyan and Timmers, 2019; Muller and Tora, 2014; Vermeulen et al., 2007) or via interaction of TF such as NF-Y (Frontini et al., 2002) that we also find to be a major TFBS enriched at promoters, consistent with previous observations (Oldfield et al., 2019).

Our data are consistent with a role of G4s in favouring pause release by Pol II. First, modelling transcription at the Eef1a1 G4 mutant promoter indicates one additional limiting step for transcription (OFF2) likely corresponding to chromatin opening in the absence of G4. Another limiting step includes transition to productive elongation, which comprises pausing, also show a significant time increase (k3, Figure 6H). Second, stabilizing G4s globally with PDS *in vivo* resulted in pause release of genes containing weaker G4s in their promoter (Figure 7C). These results are in apparent contrast with recent work indicating that treatment with another G4 ligand reduces transcription initiation (Li et al., 2021). However, the time of treatments performed in this study were much longer, opening the possibility that more indirect effects might have come into play and also allowing accumulation of DNA breaks. Overall, our data are in agreement with previous work having shown that G4-containing promoters tend to show less poised Pol II (Dao et al., 2016). How could thus G4s facilitate pause release? Because G4s at promoter exclude nucleosome, the presence of one or multiple G4s would favour not only open chromatin but also a pre-melted template for Pol II. Such structures would potentially facilitate the formation and the extension of the transcription bubble. As a consequence, crossing the +1 nucleosomal barrier would become easier for the Pol II complex, resulting in pause release. The comparison of experimental G4 signal (G4access) to ssDNA scored by KMnO4 footprinting supports this hypothesis since it indicates a correlation between G4 formation and open complex at active promoters (Figure S2E).

Another striking property of G4 forming sequences, also visible at experimental G4s determined by G4access, is their ability to position nucleosomes. The nucleosome repeat length observed is in the range of that described for CTCF but also of strongly positioning nucleosome *in vitro* (Valouev et al., 2011). Because G4s tend to be present as clusters in promoters, at these locations, nucleosome positioning is less visible probably due to the presence of multiple G4 introducing fuzziness in the adjacent nucleosomes and to their additive effects. We do not know at this stage if the positioning property in IGRs is inherent to G4 structure or could be explained by the recruitment of G4 binders. In any of these scenarios, the barrier constituted by the G4s would dock the surrounding nucleosomes. It will thus be interesting in future studies to investigate the precise interplay between G4 and CTCF since recent work suggest they could be locally associated (Tikhonova et al., 2021).

Our data point to a role of G4 as driver of CGI’s properties, possibly because they yield a more robust and/or constitutive NDR. Over 70% of G4access or G4 ChIP (Mao et al., 2018) enriched areas are actually located in CGIs and consistently, CGIs also contain large G4 clusters, increasing the likelihood of their formation locally. Our analyses suggest that pG4s could be one essential determinant of the ability of CGIs to exclude nucleosomes, thus adding a novel determinant, besides GC and CpG content (Deaton and Bird, 2011), of these essential areas of the genome. Importantly, while our study shows that G4 forming sequences behave as promoter elements by excluding nucleosome, it does not demonstrate per se that the G4 structures are formed in situ, in the context of chromatin. Nevertheless, the use of various orthogonal techniques to score for pG4s based on different principles and the pG4 ability to exclude nucleosome intrinsically pleads for their direct structural involvement as promoter element rather than protein docking sites on DNA. All in all, our work opens a new gate in our understanding and definition of a promoter *in vivo* and readjusts the existing paradigms. It will also support future work on targeting secondary structures to control their activity using specific ligands in cancer therapy.

## Methods

### Cell lines and culture

Data presented in this article were issued from the analysis of human cell lines (K562, Raji, HeLa) or mouse primary thymocytes (CD+ CD8+ (DP)). Original data presented concern essentially Raji and HeLa cells but all cellular models are described in this section.

K562 is a pseudotriploid ENCODE Tier I erythroleukemia cell line derived from a female (age 53) with chronic myelogenous leukemia. The Raji cells are lymphoblast-like cells from a male (age 11) with Burkitt’s lymphoma. HeLa Flp-in H9 cells (a kind gift of S. Emiliani) is a cell line derived from the parent HeLa line. The Hela line is derived from a female (age 31) with adenocarcinoma. Mouse CD4+ CD8+ DP cells were sorted from thymuses of 5 to 6 weeks old mice as described (Fenouil et al., 2012; Koch et al., 2011).

Raji cells were grown in RPMI 1640 medium supplemented with 10% fetal calf serum, penicillin/streptomycin (100 units/L) and glutamin (2 mg/L) at 37°C and 5% CO2. For the a-amanitin experiments (ED Figure 6A), cells were treated with 2.5 μg/L at the indicated times as described(Fenouil et al., 2012). For the PDS experiments (Figure 7 and Figure S7), cells were treated with 10 μM PDS at the indicated times.

HeLa Flp-in H9 cells used for reporter assays (Figure 5–6, Figure S6) were maintained in DMEM supplemented with 10% fetal calf serum, penicillin/streptomycin (100 units/L) and glutamin (2.9 mg/L) at 37°C and 5% CO2. HeLa cells with integrated constructs were transfected with plasmids using JetPrime (Polyplus), following manufacturer recommendations.

### Genome-wide data sets

All data sets used in this study including published and original experiments are described in Table S1. All GEO accession numbers are included. The GEO accessions for specific experiments related to this study are recorded under GSE52914.

### MNase and MNase-seq

For sequencing of nucleosomal DNA in Raji cells, 3.5×10^7^ cells were resuspended in 350 μl Solution I (150 mM sucrose, 80 mM KCl, 5 mM K_2_HPO_4_, 5 mM MgCl_2_, 0.5 mM CaCl_2_, 35 mM HEPES pH 7.4) and NP40 was added to a final concentration of 0.2%. Cell membranes were permeabilized for 5 min at 37°C. MNase was prepared at 50, 25, 12, 6 or 3 units in 0.5 mL of Solution II (150 mM sucrose, 50 mM Tris pH 8, 50 mM NaCl, 2 mM CaCl_2_) and incubated with 50 μL of cellular preparation, corresponding to 5×10^6^ cells, for exactly 10 min at 37°C. The reactions were stopped by adding EDTA to a final concentration of 10 mM. The cells were lysed using 1.45 mL of SDS Lysis Buffer (1% SDS, 10 mM EDTA pH 8, 50 mM Tris pH 8), with 10 min incubation at 4°C. 200μl aliquots were taken for purification and the remaining extracts were stored at −80°C. An equal volume of TE (200μl) was added to the aliquots, followed by subsequent 2h treatments with 0.2μg/mL of RNase A and Proteinase K at 37°C and 55°C, respectively. DNA was extracted by two subsequent phenol:chloroform:isoamylalcohol (25:24:1) extractions, further purified using QIAquick PCR purifications columns (Qiagen, Germany). Nucleosomal digestion was verified by running 500ng of DNA on a 1.5% agarose gel as well as on DNA high-sensitivity 2100 Bioanalyzer chips (Agilent, USA). Digestions showing 75% of mononucleomes (running at 150bp) were selected for library preparations. Fragments below 250 bp were purified with Ampure XP Beads (Beckman Coulter, USA) following manufacturer instructions. Librairies were prepared with TruSeq ChIP Library Preparation Kit (illumina) and sequenced on Hiseq 2000 or 4000 sequencers (Illumina).

Chromatin analysis by MNase treatment on HeLa cells (Figure 6C) were performed as follows, since adherent cells harvested with trypsin tend to clamp using the method described above. HeLa cells were harvested using trypsin and washed twice with ice-cold PBS. Cells were resuspended in 250 μL of ice-cold Nuclei buffer I (15 mM Tris-HCl pH7.5, 300 mM sucrose, 60 mM KCl, 15 mM NaCl, 5 mM MgCl_2_, 0.1 mM EGTA, 0.5 mM DTT, 0.1 mM PMSF, 3.6 μg/mL aprotinin) before addition of 250 μL of ice-cold Nuclei buffer II (15 mM Tris-HCl pH7.5, 300 mM sucrose, 60 mM KCl, 15 mM NaCl, 5 mM MgCl_2_, 0.1 mM EGTA, 0.5 mM DTT, 0.1 mM PMSF, 3.6 μg/mL aprotinin, 0.4% IGEPAL CA-630). Extracts were incubated 10 min on ice and layered on 1 mL Nuclei buffer III - sucrose cushion (15 mM Tris-HCl pH7.5, 1.2 M sucrose, 60 mM KCl, 15 mM NaCl, 5 mM MgCl_2_, 0.1 mM EGTA, 0.5 mM DTT, 0.1 mM PMSF, 3.6 μg/mL aprotinin). Nuclei were isolated by centrifugation at 10,000 g for 20 min at 4°C, and were resuspended in 600 μL of MNase digestion buffer (50 mM Tris-HCl pH7.5, 320 mM sucrose, 4 mM MgCl_2_, 1 mM CaCl_2_, 0.1 mM PMSF) and incubated on ice for 3 min. MNase was added for exactly 10 min at 37°C, using 50, 25, 12, 6 or 3 units of the enzyme. The reactions were then stopped by adding EDTA to a final concentration of 10 mM. 100 μL of SDS Lysis Buffer were added (1% SDS, 10 mM EDTA pH 8, 50 mM Tris pH 8) and after 10 min of incubation at 4°C, samples were processed as previously described for DNA purification. qPCR quantifications were performed by using the primers described in Table S4.

### G4access

The complete G4access procedure is described in(Garcia-Oliver et al., 2022) and the principle of the method is summarized in Figure S3a. In short, K562 cells were pelleted and rinsed twice in phosphate-buffered saline buffer (PBS). For each experiment, 5×10^6^ cells per titration point were re-suspended in 50 μL of prewarmed permeabilization buffer (150 mM of sucrose, 80 mM KCl, 5 mM KH_2_PO_4_, 5 mM MgCl_2_, 0.5 mM CaCl_2_ and 35 mM HEPES pH 7.4) supplemented with 0.2% (v/v) NP40 and incubated for 5 minutes at 37°C prior digestion. MNase digestions, were then performed by adding a volume of 500 μL of prewarmed MNase reaction buffer (150 mM sucrose, 50 mM Tris-HCl pH 8, 50 mM NaCl and 2 mM CaCl_2_) supplemented with either 3, 6, 12, 25 or 50U of MNase (Merck, 10107921001). Digestions were incubated at 37°C for 10 min and stopped on ice and by adding 11 μL of 500 mM EDTA to each reaction. Samples were then incubated 10 minutes on ice with 550 μL of SDS lysis buffer (1% (v/v) SDS, 10 mM EDTA and 50 mM Tris.HCl pH 8). Before DNA purification, 1 mL of water was added to dilute the SDS and the samples were incubated with 5 μL of RNAse A (ThermoFisher, EN0531) at 37 °C for 2 hours and with 8 μL of proteinase K (Euromedex, 09-0911) at 56 °C for 2 hours to complete the lysis. To then quality control the MNase digestions: 125 μL of each sample were cleaned-up using QIAquick PCR Purification Kit (QIAGEN, 28106) and assessed by agarose gel and Bioanalyzer. At this step, for efficient G4access, samples should present ~30% (+/-5%) of mono-nucleosomes. Importantly, this assessment should be performed on purified DNA that does not contain the subnucleosomal fraction, using a bioanalyzer equipment. The remaining of the samples was then purified by phenol-chloroform and ethanol precipitation for subsequent steps. We recommend that, when implementing this method, a wide range of MNase concentrations shall be tested in a first round of preparative experiments to narrow down the condition in which the critical fraction of 30% of mononucleosome shall be obtained. Our experiences showed this fraction is on average optimal for best G4 sequence recovery.

The 0-100 bp size-selected fragments from MNase digestions that have ~30% of mono-nucleosomes were subjected to DNA library preparation. In parallel, genomic DNA libraries were sonicated by Bioruptor^®^ Pico sonicator (Diagenode) to obtain DNA fragments of ~150 bp to be used later as reference data sets for bioinformatic analyses. Paired-end libraries were constructed using NEBNext^®^ Ultra™ II DNA Library Prep Kit for Illumina (New England Biolabs, E7645S) using a starting material of 50 ng. DNA fragments were treated with end-repair, A-tailing and ligation of Illumina-compatible adapters. Clean-up of adaptor-ligated DNA was performed by using CleanNGS beads (CNGS-0050) with a bead:DNA ratio of 2:1. The purified products were amplified with 8 cycles of PCR. Finally, samples were cleaned up with a bead:DNA ratio of 0.8:1 to remove the free sequencing adapters. Libraries were sequenced on the Illumina NextSeq-500 Sequencer using paired 50-30 bp reads. The G4access data is deposited to GEO database under GSE31755.

### ChIP-seq and ChIP qPCR

Fifty million cells were used to perform each Pol II ChIP-seq experiment. Cells were crosslinked for 10 min at 20°C with the crosslinking solution (10 mM NaCl, 0.1 mM EDTA pH 8, 0.05 mM EGTA pH 8, 5 mM HEPES pH 7.8 and 1% formaldehyde). The reaction was stopped by adding glycine to reach a final concentration of 250 mM. After 5 min of formaldehyde quenching, cells were washed twice with cold PBS and resuspended in cold 2.5mL LB1 (50 mM HEPES pH 7.5, 140 mM NaCl, 1 mM EDTA pH 8, 10% glycerol, 0.75% NP-40, 0.25% Triton X-100) at 4°C for 20 min on a rotating wheel. Nuclei were pelleted down by spinning at 1350 rcf in a refrigerated centrifuge and washed in 2.5mL LB2 (200 mM NaCl, 1 mM EDTA pH 8, 0.5 mM EGTA pH 8, 10 mM Tris pH 8) for 10 min at 4°C on a rotating wheel followed by centrifugation to collect nuclei. Nuclei were then resuspended in 1mL LB3 (1 mM EDTA pH 8, 0.5 mM EGTA pH 8, 10 mM Tris pH 8, 100 mM NaCl, 0.1% Na-Deoxycholate, 0.5% N-lauroylsarcosine) and sonicated using Bioruptor Pico (Diagenode) in 15mL tubes for 20 cycles of 30 s ON and 30 s OFF pulses in 4°C bath. All buffers (LB1, LB2 and LB3) were complemented with EDTA free Protease inhibitor cocktail (Roche), 0.2 mM PMSF and 1 μg/mL Pepstatin just before use. After sonication, Triton X-100 was added to a final concentration of 1% followed by centrifugation at 20000 rcf and 4°C for 10 min to remove particulate matter. After taking aside a 50 μl aliquot to serve as input and to analyze fragmentation, chromatin was aliquoted and snap-frozen in liquid nitrogen and stored at −80°C until use in ChIP assays. Input aliquots were mixed with an equal volume of 2X elution buffer (100 mM Tris pH 8.0, 20 mM EDTA, 2% SDS) and incubated at 65°C for 12 hours for reverse-crosslinking. An equal volume of TE buffer (10 mM Tris pH 8 and 1 mM EDTA pH 8) was added to dilute the SDS to 0.5% followed by treatment with RNase A (0.2μg/mL) at 37°C for one hour and Proteinase K (0.2 μg/L) for two hours at 55°C. DNA was isolated by phenol:chloroform:isoamylalcohol (25:24:1 pH 8) extraction followed by Qiaquick PCR Purification (QIAGEN, Germany). Purified DNA was then analyzed on a 1.5% agarose gel and on Bioanalyzer (Agilent, USA) using a High Sensitivity DNA Assay.

For Pol II ChIP, Protein-G coated Dynabeads were incubated at 4°C in blocking solution (0.5% BSA in PBS) carrying Pol II N20 (Santa-Cruz sc-899x, lot H3115) and TBP N12 (Santa-Cruz sc-204, lot LO214) specific antibodies. Sonicated chromatin (1mL) was added to pre-coated beads (250μL) and the mix was incubated overnight at 4°C on a rotating wheel. After incubation with chromatin, beads were washed 7 times with Wash buffer (50 mM HEPES pH 7.6, 500 mM LiCl, 1 mM EDTA pH 8, 1% NP-40, 0.7% Na-Deoxycholate, 1X protease inhibitor cocktail) followed by one wash with TE-NaCl buffer (10 mM Tris pH 8 and 1 mM EDTA pH 8, 50 mM NaCl) and a final wash with TE buffer (10 mM Tris pH 8 and 1 mM EDTA pH 8). Immunoprecipitated chromatin was eluted by two sequential incubations with 50 μL Elution buffer (50 mM Tris pH 8, 10 mM EDTA pH 8, 1% SDS) at 65°C for 15 min. The two eluates were pooled and incubated at 65°C for 12 hours to reverse-crosslink the chromatin followed by treatment with RNase A and Proteinase K and purification of DNA, as described above for input samples. Both input and ChIP samples were subjected to Bioanalyzer analysis to check that the major bulk of isolated DNA was in the 250 bp size range.

Samples were analyzed by qPCR (Stratagene) in HeLa cells following the manufacturer recommendations. Oligonucleotides pairs used for qPCR in this study are presented in the Table S4. For ChIP-seq experiments in Raji cells, purified DNA was quantified with Qubit DS DNA HS Assay (ThermoFisher Scientific, USA). Five ng of ChIP DNA were used to prepare sequencing libraries with Illumina ChIP Sample Library Prep Kit (Illumina, USA). After end-repair and adaptor ligation, library fragments were amplified by 12 cycles of PCR. Barcoded libraries from different samples were pooled together and sequenced on Illumina HiSeq2000 platform in paired-end sequencing runs.

### Chromatin RNA sequencing (chrRNA-seq)

Chromatin associated RNAs (ChrRNAs) were isolated from 2×10^7^ Raji cells before and after 10 min of PDS treatment (Figure 7F-G) as described previously (Nojima et al., 2015) followed by TurboDNase treatment. Purified RNAs were quantified by Qubit and quality was assessed using RNA Pico Assay kit with Bioanalyzer (Agilent Technologies, USA). chrRNA were then subjected to library preparation using True-seq stranded total RNA library prep gold kit (Ref#220599) from Illumina using 1 μg of chrRNA, 15 cycles of amplification and following manufacturer instructions (including ribo-depletion). The data is submitted to GEO database together with the manuscript(Garcia-Oliver et al., 2022) (Table S1).

### Bioinformatics

#### Motif analysis

Canonical promoter elements, *de novo* and known motifs (TRANSFAC) were analyzed across all expressed genes in K562, Raji and DP cells (Figure 1 and Figure S1). *De novo* motif discovery and known motif identification were performed using MEME and DREME (Bailey, 2011; Bailey et al., 2009) using fragments from −100 to +20 bp of experimental TSSs since this area encompasses not only the promoter but also the majority of the NDR. Enrichment of canonical promoter elements were tested using bedtools ‘intersect’ against all promoters of expressed genes (−100 to +20 bp of experimental TSSs) and against 10,000 permutations of random genomic areas of 121bp. Random controls were generated using bedtools ‘random’ using or not GC constraints. GC thresholds were determined to fit exactly the GC biased observed at promoters of expressed genes. Motifs used for this analysis (Table S2) were the BRE, the canonical or non-canonical TATAboxes as indicated in ED Figure 1. Additionally, we also used the quadparser QP1-7 ((G_n>2_N1-7)x4) and bed files generated by the G4Hunter algorithm G4H2.0 or G41.5 using a window of 25 bp as described (Bedrat et al., 2016).

#### G-quadruplex predictions

G-quadruplex predictions were performed using the G4Hunter(Bedrat et al., 2016) algorithm. Predicted G-quadruplexes (pG4s) at stringencies 1.52 and 2.0 were used throughout this study. Previous experiments have shown that these G4Hunter thresholds allow to experimentally confirm 92 and 100% of the predicted G-quadruplex structures(Bedrat et al., 2016).

#### ChIP-seq, ChIP-exo and MNase-seq analyses

All genomic experiments from this study or re-analyzed from available datasets were processed using our pipeline. Sequencing files were analysed using Bowtie2(Langmead and Salzberg, 2012) and PASHA(Fenouil et al., 2016). Raw sequencing reads were aligned to human Hg19 or mouse genome (mm9) using Bowtie2. Duplicate reads with identical coordinates (sequencing depth taken into account) to remove potential sequencing and alignment artifacts. For ChIP-seq and MNase-seq (nucleosome density) signal analyses, aligned reads were elongated *in silico* using the DNA fragment size inferred from paired-reads or an estimated optimal fragment size for orphan reads using Pasha R package. These elongated reads were then used to calculate the number of fragments that overlapped at a given nucleotide thus representing an enrichment score for each bin in the genome. For nucleosome positioning analyses (midpoints) presented in Figure 3, to determine the average nucleosome positions, wiggle files representing the central nucleotides of DNA fragments were also generated. For ChIP-exo, the nucleotide located at the 5’ extremity of the DNA fragments was considered to generate wiggle files, since it represents the exact points where the nucleases have stopped. Wiggle files representing average enrichment score every 50bp or 10bp were generated. Sequencing data from Input samples were treated in the same way to generate Input wiggle files. All wiggle files were then rescaled to normalize the enrichment scores to reads per million. For ChIP-seq datasets, enrichment scores from input sample wiggle files were subtracted from ChIP sample wiggle files. This allows removing/reducing the over-representation of certain genomic regions due to biased sonication, local duplications, and DNA sequencing. Finally, for MNase-seq, we smoothed the signal by replacing each 10bp bin by the average of the 5 surrounding bins on each side.

#### RNA-seq analysis

All RNA-seq datasets re-analyzed in this study were processed using our in house pipeline. Raw sequencing reads were aligned to mouse genome (mm9) or human genome (hg19) using TopHat2(Kim et al., 2013). Alignment files were then treated using PASHA(Fenouil et al., 2016) to generate wiggle files. In Raji and DP cells, experimental TSSs were determined as the summit in short-RNA-seq signals in a window of 300 bp of annotated TSSs.

#### Average binding profiles and heatmaps

To generate average binding profiles (Figure 1–4, and Figure S2-5), R scripts were developed and used for retrieving bin scores in defined regions from 10 or 50 bp bin sized wiggle files(Fenouil et al., 2016). Heatmaps were generated, viewed and color-scaled according to sample read depth using Java TreeView(Saldanha, 2004). Regions were defined either as centered on experimental TSSs (see above), on the center of predicted G4 from G4Hunter or the center of the area if no G4 was predicted. In addition, pG4s that where not located in annotated gene features or further than 200 bp from annotated TSSs were considered as intergenic.

To generate average binding profiles of Pol II and of chrRNA (Figure 7D-G), hg19 Refseq genes annotations were used to extract values from wiggle files associated with the selected genes. Bin scores inside these annotations and in a region of 5kb before the TSSs and after 5kb of annotated termination sites were determined. Based on the gene list selections, bin scores from wiggle files were used to re-scale values between TSSs and transcription termination sites (gene body) of all genes using linear interpolation. In total, 1000 points were interpolated for the gene body of each selected gene in all average profiles presented.

#### Identification of inactive promoters

To identify inactive promoters, we selected the bottom 30% of genes of Pol II signal over the defined areas (Figure 2C, Figure S2I). Hg19 Refseq genes annotations were used to extract values from wiggle files associated with the selected genes and bin scores in a region of 2kb before and after the TSSs were determined.

#### Nucleosome arrays

We have performed a precise assessment of nucleosome repeat length (NRL) and phasing, comparing pG4s and CTCF(Maurano et al., 2015) in K562 (Figure 3C), which yielded an average NRL of 194 and 193 nt calculated over 5 nucleosomes from the docking site and 215 and 216 nt over 10 nucleosomes, respectively.

#### Pausing scores

To analyze how the G-quadruplex ligand pyridostatin (PDS) impacts Pol II pausing (Figure 7B and Figure S7B), we have determined pausing scores based on the ratio of Pol II signals at promoters and in gene bodies(Adelman and Lis, 2012). Our approach for pausing scores determination is comparable to the one previously described(Fenouil et al., 2012) with modifications. It takes into account Pol II density on either promoter regions (TSS) or gene bodies (GB). Promoter regions were considered between −300 and +100 of TSS to define paused Pol II density for calculations. Densities at genes bodies were analyzed in the intervals of 50-100% of the length. The use of these intervals avoids detecting signal originating from the promoters for short genes or genes with exceptionally large initiation areas and allows detecting more significant signal of elongating Pol II. To avoid interferences between promoter and gene body read counts, only genes larger than 3kb were considered. Read count was performed using HTseq(Anders et al., 2015), normalized to the length of the genomic regions and expressed as RPKM (reads per kb per millions). Only genes with sufficient read coverage were considered (>75 RPKM at promoters and >25 RPKM at gene bodies n = 7617). Pausing scores were expressed as the ratio TSS/GB. To define a high confidence set of genes with pause release effect and since in our datasets PDS globally affected Pol II pausing (ED Figure 9b, linear regression slope T0 versus T10 minutes = 0.71, Wilcoxon test <0.00001, n= 7617), we further selected genes with significant Pol II signal increase in their gene bodies using DESEQ (Pvalue < 0.05, n= 556; Pvalue ≥ 0.05, n=7061).

### Plasmids and cloning

The repeats of the 256xMS2 binding sites were cloned from chemically-synthesized oligonucleotides into pMK123(Alexander et al., 2010). The MS2 stem loops are separated by a linker of only three nucleotides and cloned in a pIntro-MS2×256 plasmid, which also contained an FRT-Hygro cassette for Flp-in recombination(Boireau et al., 2007; Tantale et al., 2016a). pUC57 containing the Eef1a1 (1513bp) or Polr2a (711bp) WT mouse promoters were purchased at genescript; pGL4.17 containing the Pkm (200bp), Klf6 (400bp) or Taok1 (300bp) or mutagenized mouse promoters were purchased at genecust (Table S5). All construct were then subcloned into pIntro-MS2×256 between SnaBI and MluI sites. To introduce mutations into the largest promoters Polr2a and Eef1a1, smaller fragments (206 and 219 bp respectively) were purchased and subcloned to the full-length promoter between NotI and Nhe1 sites for Eef1a1 and between NotI and MluI for Polr2a. Additionally, for a second version of Eef1a1 and Polr2a WT promoters and for Eef1a1 *inv*, Polr2a*G4mut*2 and for Polr2a*G4mut3* promoters an additional luciferase reporter was added downstream of the MS2 reporter using NEB assembly builder kit and the two following oligonucleotides Gibintro-BsrG1-ires-fwd: GGTTTTCCAGTCACACCTCATGTACAGGCCCCTCTCCCTCCCCCCC and Gibluc-BsiW1-intro-Rev: TGTAAGTCATTGGTCTTAAACGTACGTCTAGAATTACACGGCGATC.

Stable expression of MCP-GFP was achieved by retroviral-mediated integration of a self-inactivating vector containing an internal ubiquitin promoter. The MCP used dimerizes in solution and contained the deltaFG deletion, the V29I mutation, and an SV40 NLS24(Tantale et al., 2016a). MCP-GFP expressing cells were grown as pool of clones and FACS-sorted to select cells expressing low levels of fluorescence. Isogenic stable cell lines expressing the reporter genes were created using the Flp-In system and a HeLa Flp-in H9 integrants were selected on hygromycin (150 μg/L). For each construct, several individual clones were picked and analysed by *in situ* hybridization. Clones usually looked similar, and two of them were further selected for the experiments after PCR and sequencing check.

### Circular dichroism (CD) spectroscopy

CD spectra were recorded on a Jasco J-815 spectropolarimeter equipped with a Peltier temperature control accessory (JASCO Co., Ltd., Hachioji, Japan). Each spectrum was obtained by averaging three scans at a speed of 100 nm/min. A background CD spectrum of corresponding buffer solution was subtracted from the average scan for each sample. The CD profile was monitored between 220 nm and 300 nm using quartz cells of 5 mm path-length and a volume of 1000 μl.

### Absorbance spectroscopy

All spectra were recorded on a Cary-300 (Agilent Technologies) spectrophotometer in 10 mM lithium cacodylate buffer (pH 7.2) at 3 or 4 μM oligonucleotide strand concentration, in the presence or absence of 100 mM KCl.

Thermal difference spectra (TDS) were obtained by taking the difference between the absorbance spectra of unfolded and folded oligonucleotides that were recorded at high (95°C) and low (25°C) temperatures, respectively, in a buffer containing 100 mM KCl. TDS provides specific signatures of different DNA structural conformations, provided that the structure is not too heat-stable (a number of G4 structures do not melt at high temperatures).

Isothermal difference spectra (IDS) were obtained as described previously(Renaud de la Faverie et al., 2014) by taking the difference between the absorbance spectra from unfolded and folded oligonucleotides. These spectra were recorded at 25°C before and after potassium cation addition (100 mM KCl), respectively. IDS provide specific signatures of different DNA structural conformations.

### Acquisition and analysis of smFISH images

SmFISH was performed as previously described (Tantale et al., 2016a), with a mix of 10 fluorescent oligos hybridizing against the MS2×32 repeat, each oligo containing four molecules of Cy3. Since each oligo bound eight times across the MS2×256 repeats, each molecule of pre-mRNA hybridized with 80 oligos, thereby providing excellent single molecule detection and signal-to-noise ratios.

To obtain the number of released, nucleoplasmic and nascent mRNA per cell, smFISH images were recorded with an upright widefield Leica microscope as 3D image stacks with a z-spacing of 0.3 μM, with a x100 objective, and an Evolve 512×512 EMCCD camera (Photometrics). The images were analyzed with FISH-quant (Mueller et al., 2013) to count the number of pre-mRNA per nuclei, using populations of 400–500 cells per experiment. To obtain the number of nascent pre-mRNA per cell, the transcription sites (TS) were identified manually and isolated pre-mRNA molecules located in the nucleoplasm were used to define the point spread function (PSF) and the total light intensity of single molecules, which finally allowed determining the intensity of TS expressed in number of full-length transcripts.

### Live-cell image acquisition

Cells were plated on 25-mm diameter coverslips (0.17-mm thick). After 24-48 hours the coverslips were mounted in the GFP-imaging medium (DMEM-GFP-2, Evrogen) with rutin in a temperature-controlled chamber with CO_2_ and imaged on an inverted OMX Deltavision microscope in time-lapse mode. A x100, NA 1.4 objective was used, with an intermediate x2 lens and an Evolve 512×512 EMCCD camera (Photometrics). Stacks of 11 planes with a z-spacing of 0.6 μm were acquired, with one stack collected every 3 min for 8h.

### Quantification of short movies

Short movies were analysed as previously described (Tantale et al., 2021; Tantale et al., 2016a) (Figure 6G). In short, we manually defined the nuclear outline and the region within which the transcription site (TS) is visible and stacks were corrected for photobleaching using a fitted curve with a sum of three exponentials. This curve was used to normalize each time-point such as nuclear intensities were equal to the intensity of the first time-point. We then filtered the image with a 2-state Gaussian filter. First, the image was convolved with a larger kernel to obtain a background image, which was then subtracted from the original image before the quantification is performed. Second, the background-subtracted image was smoothened with a smaller Kernel, which enhances the SNR of single particles to facilitate spot pre-detection. TS positions in each frame of the filtered images were determined as the brightest pixel above a user-defined threshold in the pre-detected region of the TS. When no pixel was above the threshold, the last known TS position was used. Then the TS signal was fitted with a 3D Gaussian estimating its standard deviation *σ_×z_* and *σ∑*, amplitude, background, and position. We performed two rounds of fitting: in the first round all fitting parameters were unconstrained. In the second round, the allowed range was restricted for some parameters, to reduce large fluctuations in the estimates especially for the frames with a dim or no detectable TS. More specifically, the *σ_×z_* and *σ_z_* were restricted to the estimated median value ± standard deviation from the frames where the TS could be pre-detected, and the background was restricted to the median value. The TS intensity was finally quantified by estimating the integrated intensity above background expressed in arbitrary intensity units. With the live cell acquisition settings, the illumination power was low and we could not reliably detect all individual molecules. We therefore collected right after the end of the movies one 3D stack with increased laser intensity (50% of max intensity, compared to 1% for the movie), which allowed reliable detection of individual RNA molecules. We also collected slices with a smaller z-spacing for a better quantification accuracy (21 slices every 300 nm). Quantification of TS site intensity in the calibration stack was done with *FISH-quant* as follows: (a) when calculating the averaged image of single RNA molecules, we subtracted the estimated background from each cell to minimize the impact of the different backgrounds; (b) when quantifying the TS in a given cell, we rescaled the average image of single RNA molecules such that it had the same integrated intensity as the molecules detected in the analyzed cell. To calibrate the TS intensities in the entire movie, i.e. to express the TS intensity as a number of equivalent full-length transcripts, we used the fact that the last movie frame was acquired at the same time as the calibration stack. We then normalized the extracted TS intensity in the movies, *I*_MS2_, to get the nascent counts *N*_nasc;calib_: *N*_nasc;calib_(*t*)=*IMS*2(*t*)×(*N*hasc,final/*I*_final_), where *M*_nasc,final_ stands for the estimated number of nascent transcripts in the calibration stack and *I*_final_ for the averaged intensity of the last four frames.

### Analysis of long movies

To quantify long movies acquired at low frames rate (one 3D image stack every 3min), we used ON-quant, a rapid analysis tool that identifies transcription sites, measures their intensities, attributes the ON or OFF states of transcription, based on the defined intensity threshold under which a TS is considered to be silent, and above which a TS is considered to be active. The intensity threshold was defined based on the mean intensity of single molecules (Tantale et al., 2016a).

### Mathematical modelling, short and long movies analysis

Intermittent transcriptional activity of the promoters is modelled using a Markov process with one active and multiple inactive epigenetic states. The number of states and the transition rate parameters are obtained using the algorithms and pipeline first described in (Tantale et al., 2021).

For the sake of consistency, we provide a short description of the pipeline. The cells were imaged live for 30 minutes every 3 seconds (short movies), or for 8-9h every 3 minutes (long movies).

#### Deconvolution and position of Pol II initiation events in short movies

The Pol II positions were found by combining a genetic algorithm with a local optimisation procedure. Before initiation of the analysis algorithm, several key parameters were established. The Pol II elongation speed was fixed at 67 bp/s(Tantale et al., 2021; Tantale et al., 2016b). The reporter construct transcript was divided into three sections consisting of the pre-MS2 fragment (PRE=700 bp), 256xMS2 loops (SEQ=5800 bp), and post-MS2 fragment until the polyA site (POST=1600 bp). An extra time *P_poly_*=100s was added to POST, corresponding to the polyadenylation signal (during this time the polymerase is past the polyA site and remains on the DNA, see(Tantale et al., 2016b)). The frame rate of short movies is sufficient to detect processes that occur on the order of seconds.

In order to find the positions of initiation events via the deconvolution pipeline, all the possible initiation times were discretized using a step size of 0.45 seconds (or 30 bp at 67 bp/s). This step was chosen as it is smaller than the minimum polymerase spacing and large enough to still accommodate a reasonable computation time. For a movie of 30 min length this choice corresponds to a maximum number of 4020 positions. The deconvolution algorithm was implemented in Matlab R2020a using Global Optimization and Parallel Computing Toolboxes for optimizing Pol II positions in parallel for all nuclei in a collection of movies. Waiting times were then computed from the position of each initiation event (Figure 6G).

#### Long movies waiting time distribution

For long movies, the low resolution (3 min) does not allow a precise positioning of initiation events. In this case we binarize the signal by considering that the transcription site is active or inactive if the measured intensity is above or below a threshold level, respectively, which is set to be slightly higher than the intensity similar of a single polymerase The inactive intervals then indicate long waiting times between successive polymerases. The active intervals are used to estimate the probability that waiting times are larger than the movie frame rate (3 min), which is one of the parameters needed for connecting long and short time distributions and obtain a multiscale distribution (see (Tantale et al., 2021)).

#### Multi-exponential regression fitting of the survival function and model reverse engineering using the survival function

Waiting times were extracted as differences between successive Pol II initiation events from all the resulting traces and the corresponding data was used to estimate the nonparametric cumulative short movie distribution function by the Meyer-Kaplan method. Data from long movies is used to generate the nonparametric cumulative long movie distribution function. The two distribution functions are fitted together into a multiscale cumulative distribution function using the total probability theorem and estimates of two parameters *p_l_* and *p_s_*, representing the probabilities that waiting times are longer than the long movie frame rate, and longer than the length of the short movie, respectively (see ref(Tantale et al., 2021) for details).

Then, a multi-exponential regression fitting of the multiscale distribution function produces a set of 2N-1 distribution parameters, where N is the number of exponentials in the regression procedure (3 for N=2 and 5 for N=3). The regression procedure was initiated with multiple log-uniformly distributed initial guesses and followed by local gradient optimisation. It resulted in a best-fit solution with additional suboptimal solutions (local optima with objective function value larger than the best fit).

The 2N-1 distribution parameters can be computed from the 2N-1 kinetic parameters of a N state transcriptional bursting model. Conversely, a symbolic solution for the inverse problem was obtained, allowing computation of the kinetic parameters from the distribution parameters and reverse engineering of the transcriptional bursting model. In particular, it is possible to know exactly when the inverse problem is well-posed, i.e. when there is a unique solution in terms of kinetic parameters for any given distribution parameters in a domain (Figure 6H and Figure S6I).

#### Transcriptional bursting models

The transcriptional bursting models used in this paper are as following:

For a promoter two-state model (N=2), the model corresponds to the well-known ON-OFF telegraph model. In this case there are 3 distribution parameters and 3 transition rates parameters.

The distribution parameters are *A*_1_, *λ*_1_, *λ*_2_ defining the survival function

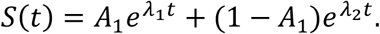

These parameters are obtained by bi-exponential fit of the empirical survival function.

The transition rates parameters of the ON-OFF telegraph model can be obtained from the distribution parameters using the formulas

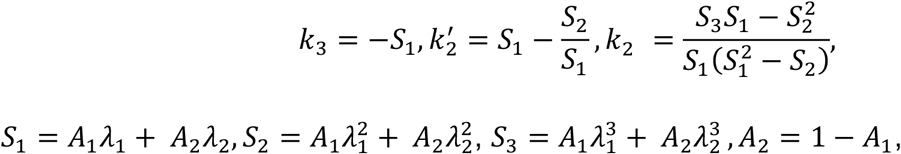

where 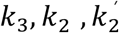 are the initiation rate, the OFF to ON and ON to OFF transition rates, respectively.

For a promoter three-state model (N=3), there are 5 distribution parameters and 5 kinetic parameters.

The distribution parameters are *A*_1_, *A*_2_, *λ*_1_, *λ*_2_, *λ*_3_, defining the survival function

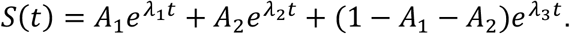

The corresponding model represented in the Figure has two OFF and one ON state. The five transition rate parameters can be obtained from the distribution parameters:

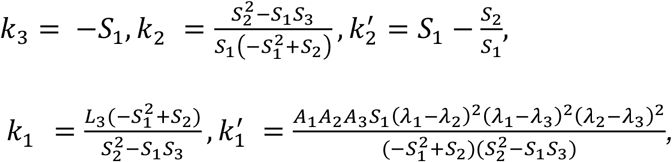

where

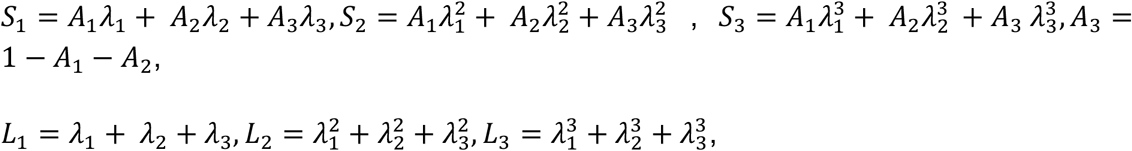

and 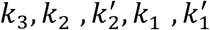 are the transcription initiation, OFF2 to ON, ON to OFF2, OFF1 to OFF2, and OFF2 to OFF1 rates, respectively.

#### Error intervals

Distribution parameters result from multi-exponential regression fitting using gradient methods with multiple initial data. These optimization methods provide a best fit (global optimum) but also suboptimal parameter values. Using an overflow ratio (a number larger than one, in our case 2) to restrict the number of suboptimal solutions, we define boundaries of the error interval as the minimum and maximum parameter values compatible with an objective function less than the best fit times the overflow.

#### Choice of the number of exponentials

The number of exponentials was determined by a parsimony principle: we have chosen the smallest N that fits well. More precisely, starting with N=2, we have increased N as long as the goodness of fit reduced without increase of overfitting. We have used parametric uncertainty (error intervals) as a proxy for overfitting (Figure S6H).

## Code availability

Softwares and codes are all publicly available and have been previously described (Descostes et al., 2014; Fenouil et al., 2012; Fenouil et al., 2016).

## Data availability

The GEO accessions for specific experiments related to this study are recorded under GSE52914.

## Acknowledgements

In the JCA lab, this work was supported by institutional grants from the CNRS including an 80prime2021 ‘Deciph G4’, and a grant from “Agence Nationale de la Recherche” (G4access, ANR-20-CE12-0023), ‘amorçage jeunes équipes’ Fondation pour la Recherche Medicale FRM AJE20130728183 and INCA PLBIO20-225. CE was supported by a grant from ARC (retour postdoc), EGO by a grant from EpiGenMed labex of excellence. JLM was supported by Inserm, CNRS and Ecole Polytechnique. We are grateful to Salvatore Spicuglia, Dan Fisher, Robert Feil and Mounia Lagha for critical reading of the manuscript and to Florian Mueller for help with quantification of live cells imaging data.

## Author contributions

JCA and CE designed experiments; CE, EGO, TM, KG and AP performed the experiments; CE and AZEA analyzed genomic data; CE, EB, MK, MCR and EB performed and interpreted the microscopy experiments, YL, AC, DV and JLM characterized and interpreted the G-quadruplexes *in vitro*, RM performed the eSNP analysis. OR performed the mathematical modelling. JCA and CE conceived the project, suggested and interpreted experiments, and wrote the article. All authors reviewed the manuscript.

## Competing interests

Authors declare no competing interests.

## Supplementary data

**Figure S1: Analysis of promoter elements and motifs in 3 independent cell lines.**

(a) Definition of major TSS (mTSS), MEME and DREME promoter motif analyses in K562 cells. CAGE datasets (FANTOM consortium) were used to define mTSSs at the nucleotide resolution (n=8346). Heatmaps show short chromatin (nascent) RNA-seq (analysed from GSE52914) and Pol II ChIP-seq (ENCODE) docked on the main sense mTSS and ranked by increasing distance between sense and antisense short RNAs (top left panel). Sequence motif analysis of the transcription initiating nucleotides (INR) at the sense and antisense mTSSs are shown (bottom left). On the right panels are shown TFBS analyses using MEME or DREME and frequency analyses for all sequences features including Quadparser (QP1-7) and BRE motifs. The random control column depicts a search for the motif in 8346 random genomic sequences (with 10000 permutations). See also Table S2 for detailed frequencies of all motifs.

(b) Definition of mTSSs and MEME and DREME motif analyses in Raji B cells. The analysis was performed as in (a) over 8356 human promoters and using chrRNA-seq in Raji for mTSS determination (analysed from GSE52914).

(c) Definition of mTSSs, MEME and DREME motif analyses in mouse primary T cells. The analysis was performed as in (a) over 7947 mouse promoters and using short-RNA datasets (size-selected below 50 bp, analysed from GSE38577) to define mTSSs at the nucleotide resolution. Short RNA-seq and Pol II ChIP-seq data sets are shown.

**Figure S2: Association of the TATAbox and G4 motifs with transcription initiation**

(a) pG4, BRE and SP1 motifs largely overlap at active promoters. Venn diagram of active promoters containing SP1, BRE motifs or pG4s. G4Hunter is displayed at two stringencies (1.5, red circle or 2.0, dotted inner circle). The principle of the G4Hunter algorithm is to score positively Gs and G stretches, while scoring negatively Cs and C stretches within a defined window (typically 25nt). Overlapping G4s, above a defined threshold, are concatenated. A and T nucleotides score are fixed as null. G4Hunter scores >1.5 and 2.0 correspond to likelihood of G4 formation *in vitro* of >95 and 99% (Bedrat et al., 2016; Garcia-Oliver et al., 2022).

(b) Promoters containing G4 predictions (G4H1.5 or 2.0) tend to harbour less other TFBS or promoter elements as compared to all promoters. This analysis was performed with the selection described in Figure 1 for K562 cells.

(c) Promoters with TATA boxes show more focused and directional transcription. Pol II ChIP-seq profiles in K562 cells (ENCODE) at promoters of expressed genes that contain either no TATA, a non-canonical (TATAW) or a canonical (TATAWAAG) TATAbox.

(d) Experimental G4 signals definition. Groups 1-6 correspond to increasing level of G4access signals. Heatmaps in K562 of predicted G4 (H4hunter 2.0) and G4 signals using G4access (Garcia-Oliver et al., 2022), G4-ChIP (Hansel-Hertsch et al., 2016; Mao et al., 2018) and G4seq (Chambers et al., 2015); in Raji of predicted G4 (H4hunter 2.0) and G4 signals using G4access (Garcia-Oliver et al., 2022), ssDNA-seq (Kouzine et al., 2013) and G4seq (Chambers et al., 2015). See also Figure 1E and 2B. The heatmaps were centered on the G4 motif upstream of TSS as shown below the tracks.

(e) Experimental G4 signal metaprofiles. Metaprofiles of the heatmaps shown in d. The signals were divided in 6 groups of ascending G4 access signals. Metaprofiles were centered on the G4 motif upstream of TSS as shown below the tracks.

(f) Metaprofiles of R-loops at active promoters and docked on pG4s (G4H 2.0) upstream of experimental TSSs in K562 cells. The signals were divided in 6 groups of ascending G4 access signals

(g) Metaprofiles of ChIP-seq of Pol II centered on pG4s (G4H2.0) on 1444 pG4-containing promoters of active genes (−100,+20 bp) in Raji cell.

(h) Metaprofiles of ChIP-seq of Pol II, TBP, TFIIB, centered on pG4s (G4H2.0) on 1291 pG4-containing promoters of active genes (−100,+20 bp) in mouse T cells (GSE38577).

(i) Metaprofiles of Polycomb-deposited H3K27me3 mark, GC and CpG-content in groups 1 and 2 defined in Figure 2C.

A green arrow indicates the sense of transcription in each graph,

All accession numbers are presented in the table S1.

**Figure S3: pG4-dependent nucleosome exclusion does not depend on SP1 binding in K562 cells.**

(a) Heatmaps ranked by increasing signals of SP1 binding and centred on a pG4 (ChIP-seq, ENCODE). All promoters of expressed genes that contain a pG4 in K562 cells are presented (n=1766). Group 1 and 2 are depleted or enriched for SP1, respectively. Metaprofiles derived from the heatmaps (groups 1 and 2) of SP1 density, GC content nucleosome densities and G4access are shown on the right.

(b) Metaprofiles of promoters not containing a canonical GC box/SP1 binding site.

(c) Metaprofiles of promoters not containing a non-canonical GC box/SP1 binding site.

Green arrows indicate the sense of transcription in each graph.

**Figure S4: pG4s show intrinsic nucleosome eviction property**

(a) Persistence of NDRs at pG4 sites following transcription inhibition by α-amanitin. Raji cells were treated for 0, 12, 18 or 36 h with 2.5 μg/mL α-amanitin. Pol II clearance and nucleosome depletion mapped by MNase-seq from active genes containing pG4s (G4H2.0) are shown at the different time points (GSE38577 and this study). A green arrow indicates the sense of transcription in each graph

(b) Metaprofiles of MNase-seq from *in vitro* reconstituted chromatin on T cell genomic DNA (GSE25133) centred on pG4 at the active pG4-containing promoters (−100, +20 bp), intergenic or intragenic regions defined in K562 cells.

(c) Heatmaps ranked by increasing MNase signals showing nucleosome or G4H1.5 (docked on G4H2.0) signals. Six manually defined groups based on relative nucleosome densities are further plotted as graphs below the heatmaps.

**Figure S5: G4s contribute to CpG islands openness and activity.**

**(a)** Selection of G4-containg and G4-depleted CpG islands. The selections were perfomed on the same number of sequences (2 x 1112) with similar length and GC content, with the indicated G4Hunter thresholds. For the groups to be of equal size, equivalent length and GC content the initial populations of CGIs with G4H>1.5 (21536) or G4H<1.2 (2191) were randomized to end up with 2 groups of 1112 sequences.

**(b)** Heatmaps as in Figure S8a for the selections presented in a. The DNAse data used here indicate more chromatin opening in the G4-containing group (ENCODE data for K562 cells, GSE32970).

**(c)** Average profiles of the group 1 and 2 shown in a and b. For Nucleosome density, a zoom over 4kb is indicated to best show the differences in the 2 groups. The differences are observed essentially for the width of the NDRs.

**Figure S6: G4s are required for full activities of model promoters**

(a) –(e) G4access (Garcia-Oliver et al., 2022) and CUT&Tag (Lyu et al., 2022) at model promoters. Signals were extracted from data obtained in mouse ES cell lines at indicated model promoters. Predicted G4s scored by G4Hunter are indicated. Their sequences, used for promoter assays in single cells, are shown below the tracks in the zoomed areas. Each of the G tracks (n>1) are indicated in light blue. Complete sequences of the model core promoters are indicated in Table S3.

(f) Quantification of smFISH images of MS2 reporter activity of Eef1a1, WT and G4inv mutant where the strand of the canonical G4 has been inverted. Mann-Whitney test was used (ns: P > 0.05). Scheme of the promoters and mutations are indicated below the charts. pG4s are represented in red rectangles (changes in font orientation indicates swap of strand) and TATAboxes in green rectangles.

(g) Representative images of long movie analyses. Long movies of MS2 reporter activity in WT and mutant cell lines as indicated. (Left) maximum image projections of selected 3-D image stacks from the 8h movies. The arrows indicate transcription sites. (Right) graphs display the corresponding quantifications of the movies, with the bars representing ON (green) and OFF (red) states.

(h) Survival functions describing transcriptional bursting at the model promoters and deconvolution indicating 2 fits to describe optimal function in the WT situation and 3 fits for both TATA and G4 mutants.

(i) WT promoters require 2 main steps for transcription, and mutant promoters 3 main steps defined by 3 or 5 time constants, respectively. The values of the derived time constants (described in methods) are indicated in the right panel.

**Figure S7: Time course analysis of G4s stabilization by PDS effects on nucleosome exclusion and positioning, and Pol II pausing.**

(a) FACS analysis of gH2AX levels following PDS treatment at the indicated time (n=3).

(b) PDS influences Pol II pausing. Changes of Pol II pausing scores (see methods) in response to 10 μM PDS of all genes with detectable Pol II at promoters and gene bodies for 0,10, 30 and 60 min are displayed. Genes that have increased gene body signals with a Pval<0.05 at t=10 min (n=556) are highlighted in dark blue (t=10 min), dark green (t=30 min) or dark orange (t=60 min), showing that pause release is relatively transient following treatment. Changes in pausing scores, promoter and gene body signals are shown.

(c) Gene ontology analysis of the 556 selected pause release genes using DAVID webtool.

**Table S1: Genomic files**

Table summarizing all resources of the genomic analyses presented in this study. The table depicts the cell type, the experiment type and the source of the files.

**Table S2: Motif search analysis**

Frequency analyses of sequence features in a 120 bp window around experimental TSSs (−100, +20). 14 motifs or features were analysed (BREd, BREu, ETS, G4H1.5, G4H2.0, QP1-7, INR, NF-Y, Half G-quadruplex, SP1 motifs, two TATAbox consensus and two negative control motifs).

**Table S3: Sequence of all mouse model promoters used in this study.**

These sequences were inserted upstream of the MS2 reporter and inserted in Hela genome using FRT system.

**Table S4: Table of qPCR oligonucleotides used in this study**

All oligonucleotide pairs used for qPCR are presented; sequences, complementary genomes and genomic location of amplicons are described.

## Notes

### Competing Interest Statement

The authors have declared no competing interest.

