## Supplemental Figures 1 to 7 for "G-quadruplexes are promoter elements controlling nucleosome exclusion and RNA polymerase II pausing"

### Esnault *et al*., Figure S1

A

#### K562 EL cells (n=8346)

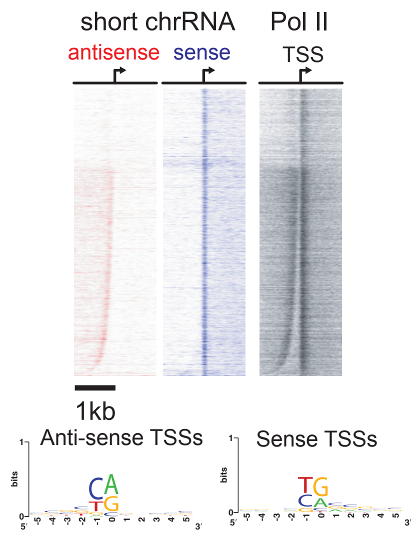


| Motif | MEME Sites | MEME E-value | MEME Rank | MEME Seq. Logo | DREME E-value | DREME Rank | DREME Seq. Logo |
|---|---|---|---|---|---|---|---|
| **G4/SP1** | 2846 | $2.2e^{-155}$ | 1 | | $1.4e^{-298}$ | 1 | |
| **Ets** | 1308 | $1.2e^{-32}$ | 2 | | $3.7e^{-194}$ | 2 | |
| **NFY** | 1071 | $2.2e^{-20}$ | 3 | | $1.2e^{-104}$ | 3 | |
| **TATA** | | n.a | | | $6.6e^{-57}$ | 9 | |

| | % in promoters | % (cor. for GC) | obs./exp. | Pvalue | Random cont. |
|---|---|---|---|---|---|
| BRE (SSRCGCC) | 45.2% (3772) | 6.3% (528) | 7.1 | $<1e^{-99}$ | 2.9% (242) |
| **G4 (G4H2.0)** | **21.1%** (1766) | 3.1% (262) | **6.7** | $<1e^{-99}$ | 1.5% (121) |
| **G4 (QP1-7)** | **21.0%** (1757) | 3.2% (274) | **6.4** | $<1e^{-99}$ | 1.5% (125) |
| **G4 (G4H1.5)** | **45.1%** (3763) | 12.0% (1005) | 3.7 | $<1e^{-99}$ | 5.9% (491) |
| TATA (TATAWAAG) | 1.0% (80) | 0.6% (46) | 1.7 | $<1e^{-99}$ | 1.5% (127) |
| TATA (TATAW) | 8.6% (721) | 16.5% (1373) | 0.5 | ns | 33.7% (2811) |

B

#### Raji B cells (n=8356)

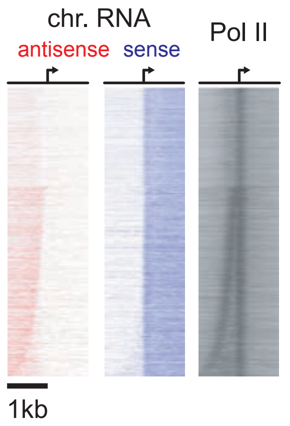


| Motif | MEME Sites | MEME E-value | MEME Rank | MEME Seq. Logo | DREME E-value | DREME Rank | DREME Seq. Logo |
|---|---|---|---|---|---|---|---|
| **G4/SP1** | 3230 | $3.9e^{-238}$ | 1 | | $7.4e^{-238}$ | 2 | |
| **Ets** | 2187 | $1.0e^{-96}$ | 2 | | $7.3e^{-260}$ | 1 | |
| **NFY** | 934 | $6.9e^{-57}$ | 3 | | $8.0e^{-156}$ | 3 | |
| **TATA** | | n.a | | | $3.5e^{-28}$ | 10 | |

| | % in promoters | % (cor. for GC) | obs./exp. | Pvalue |
|---|---|---|---|---|
| BRE (SSRCGCC) | 44.0% (3680) | 6.3% (529) | 7.0 | $<1e^{-99}$ |
| **G4 (G4H2.0)** | **17.3%** (1444) | 3.1% (262) | **5.5** | $<1e^{-99}$ |
| **G4 (QP1-7)** | **17.6%** (1467) | 3.2% (275) | **5.3** | $<1e^{-99}$ |
| **G4 (G4H1.5)** | **40.7%** (3398) | 12.0% (1007) | 3.4 | $<1e^{-99}$ |
| TATA (TATAWAAG) | 0.9% (73) | 0.6% (47) | 1.6 | $<1e^{-99}$ |
| TATA (TATAW) | 6.3% (523) | 16.5% (1375) | 0.4 | ns |

C

#### Mouse primary T cells (n=7947)

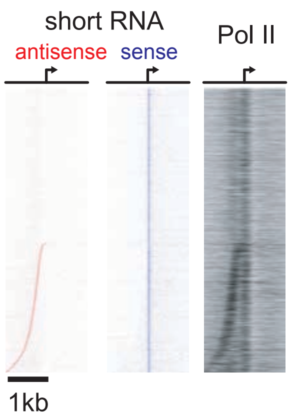


| MEME Motif | MEME Sites | MEME E-value | MEME Rank | MEME Seq. Logo | DREME Motif | DREME E-value | DREME Rank | DREME Seq. Logo |
|---|---|---|---|---|---|---|---|---|
| **G4/SP1** | 2249 | $3.9e^{-238}$ | 1 | | **G4/SP1** | $1.5e^{-227}$ | 1 | |
| **A-stretch** | 210 | $2.1e^{-47}$ | 2 | | **Ets** | $1.3e^{-135}$ | 2 | |
| **Ets or NFY** | | n.a | | | **NFY** | $1.8e^{-96}$ | 3 | |
| **TATA** | | n.a | | | **TATA** | $6.3e^{-46}$ | 7 | |

| | % in promoters | % (cor. for GC) | obs./exp. | Pvalue |
|---|---|---|---|---|
| BRE (SSRCGCC) | 42.3% (3364) | 6.5% (512) | 6.6 | $<1e^{-99}$ |
| **G4 (G4H2.0)** | **16.2%** (1291) | 3.0% (235) | **5.5** | $<1e^{-99}$ |
| **G4 (QP1-7)** | **16.3%** (1294) | 3.0% (238) | **5.4** | $<1e^{-99}$ |
| **G4 (G4H1.5)** | **40.1%** (3184) | 11.0% (944) | 3.4 | $<1e^{-99}$ |
| TATA (TATAWAAG) | 1.0% (83) | 0.6% (46) | 1.8 | $<1e^{-99}$ |
| TATA (TATAW) | 6.4% (509) | 16.9% (1345) | 0.4 | ns |

### Esnault *et al.*, Figure S2

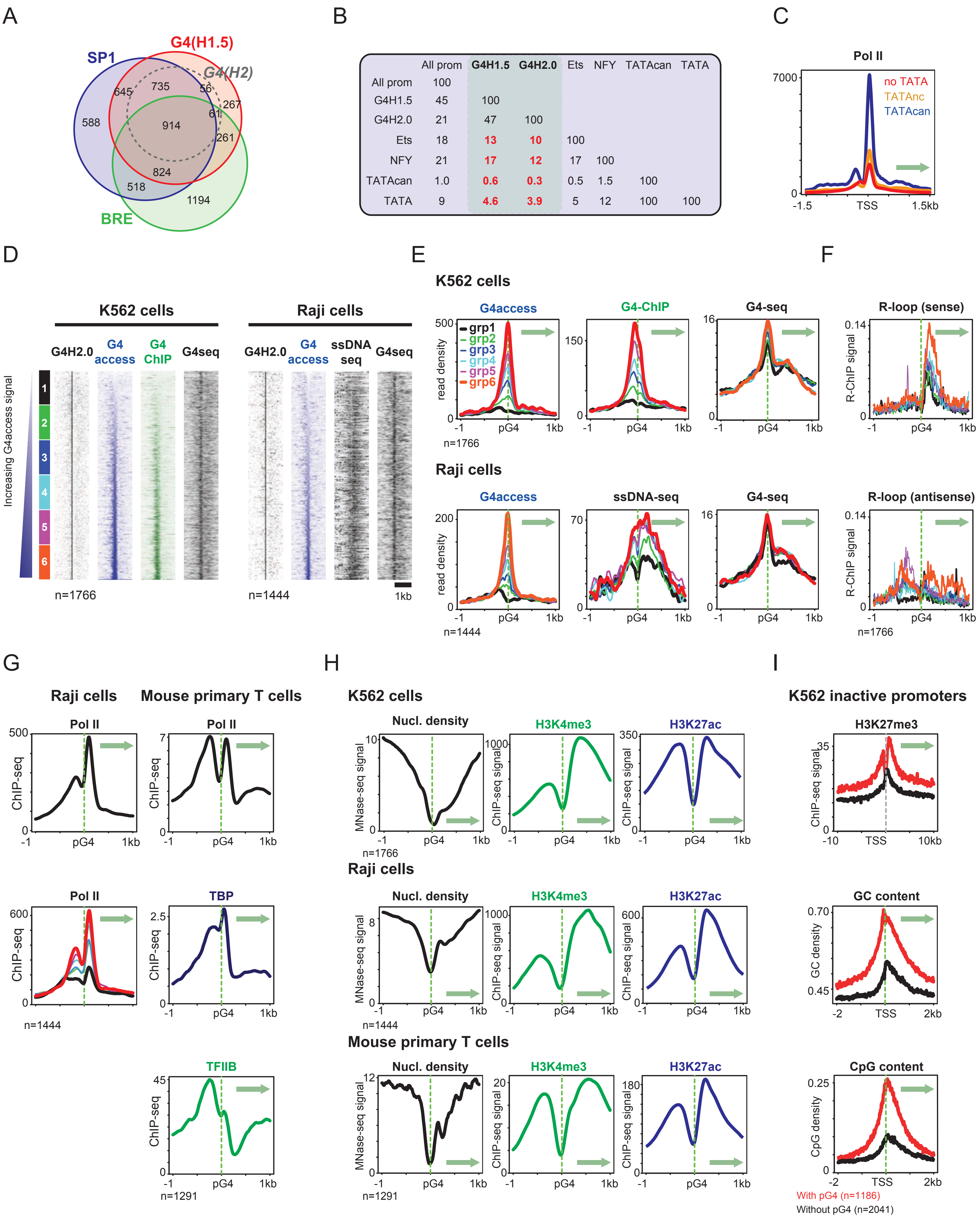


|  | All prom | G4H1.5 | G4H2.0 | Ets | NFY | TATAcan | TATA |
|---|---|---|---|---|---|---|---|
| All prom | 100 | | | | | | |
| G4H1.5 | 45 | 100 | | | | | |
| G4H2.0 | 21 | 47 | 100 | | | | |
| Ets | 18 | 13 | 10 | 100 | | | |
| NFY | 21 | 17 | 12 | 17 | 100 | | |
| TATAcan | 1.0 | 0.6 | 0.3 | 0.5 | 1.5 | 100 | |
| TATA | 9 | 4.6 | 3.9 | 5 | 12 | 100 | 100 |

**Esnault *et al.*, Figure S3**

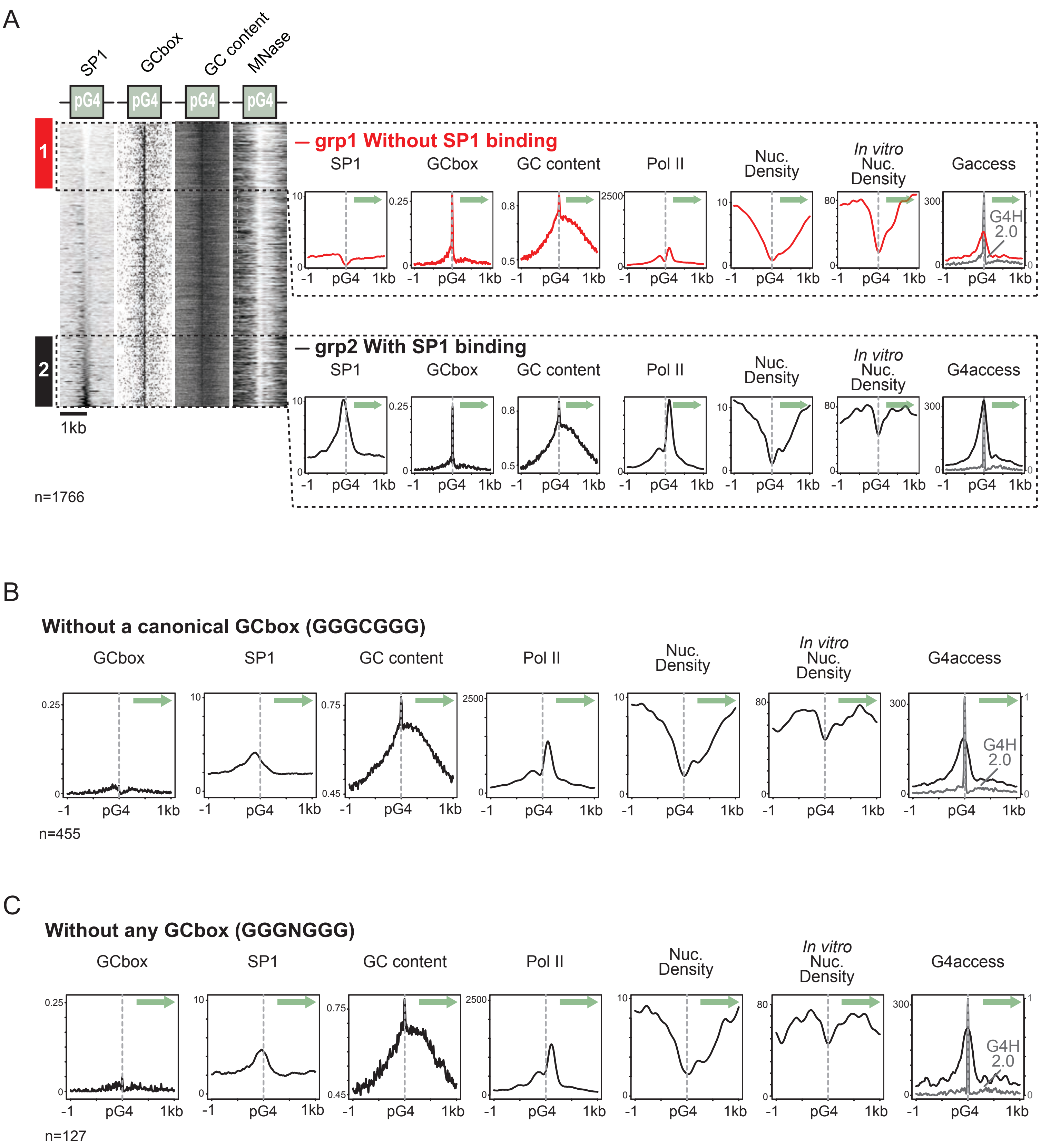


0

**Esnault *et al.*, Figure S4**

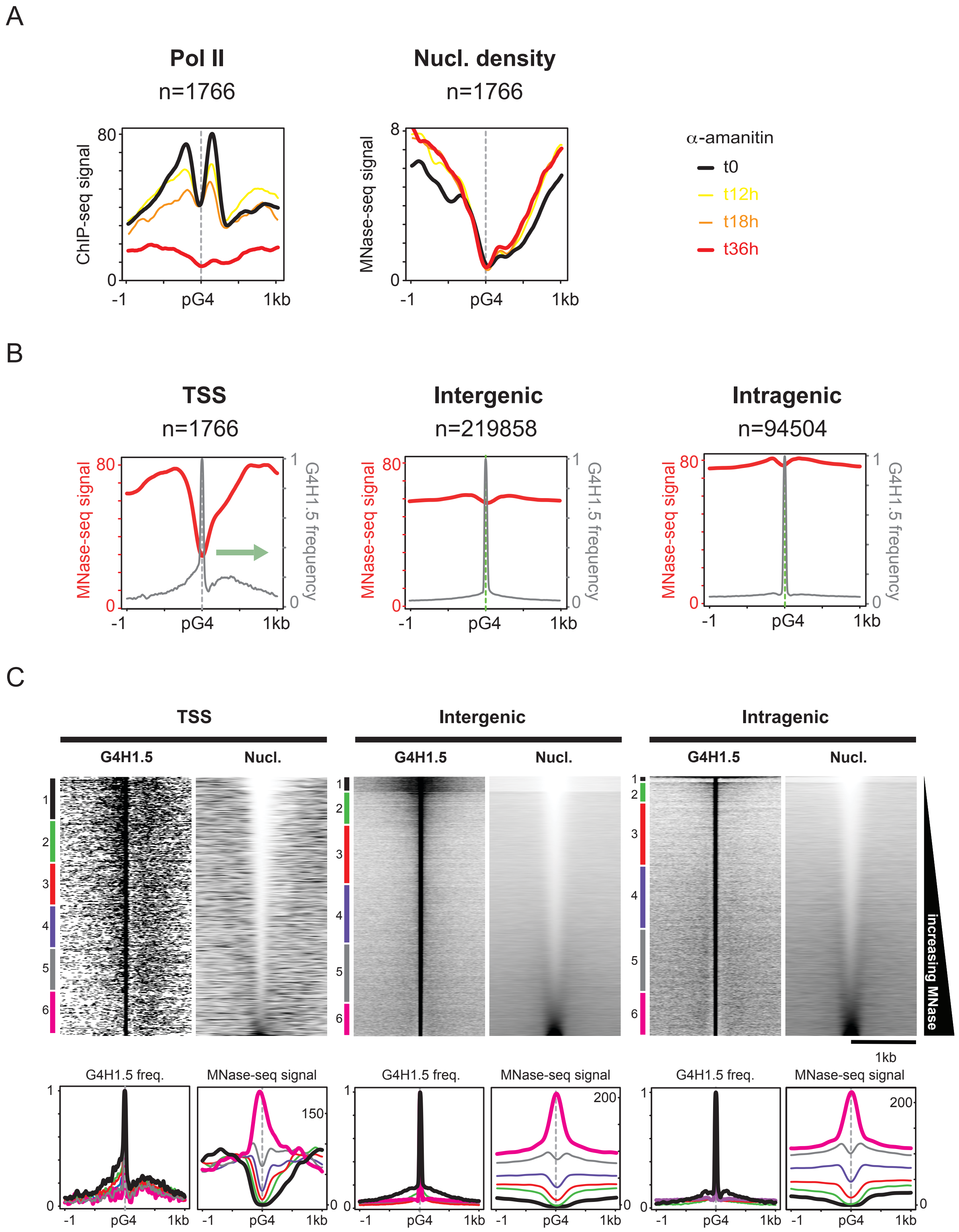

**Esnault *et al*., Figure S5**

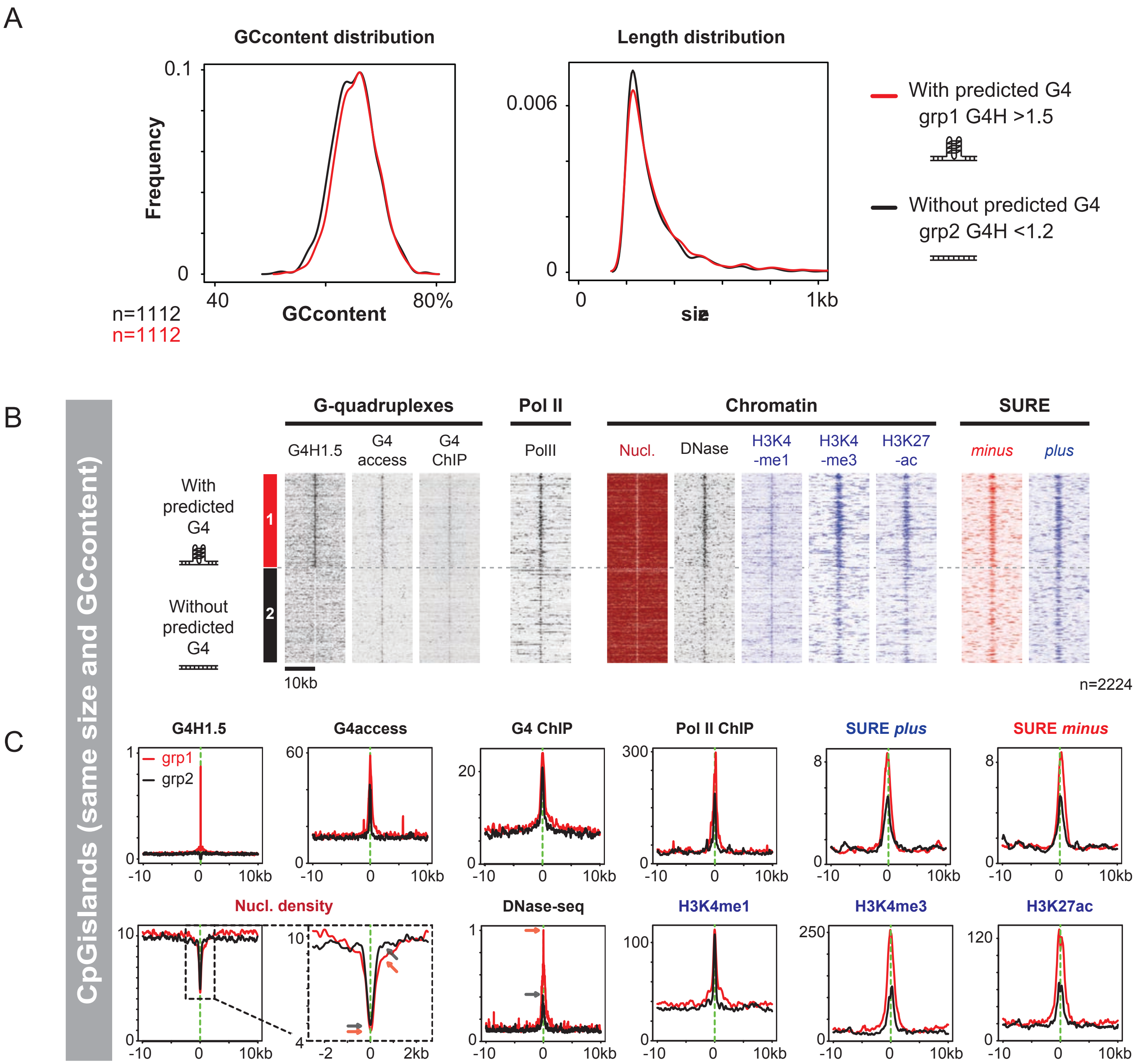

**Esnault *et al.*, Figure S6**

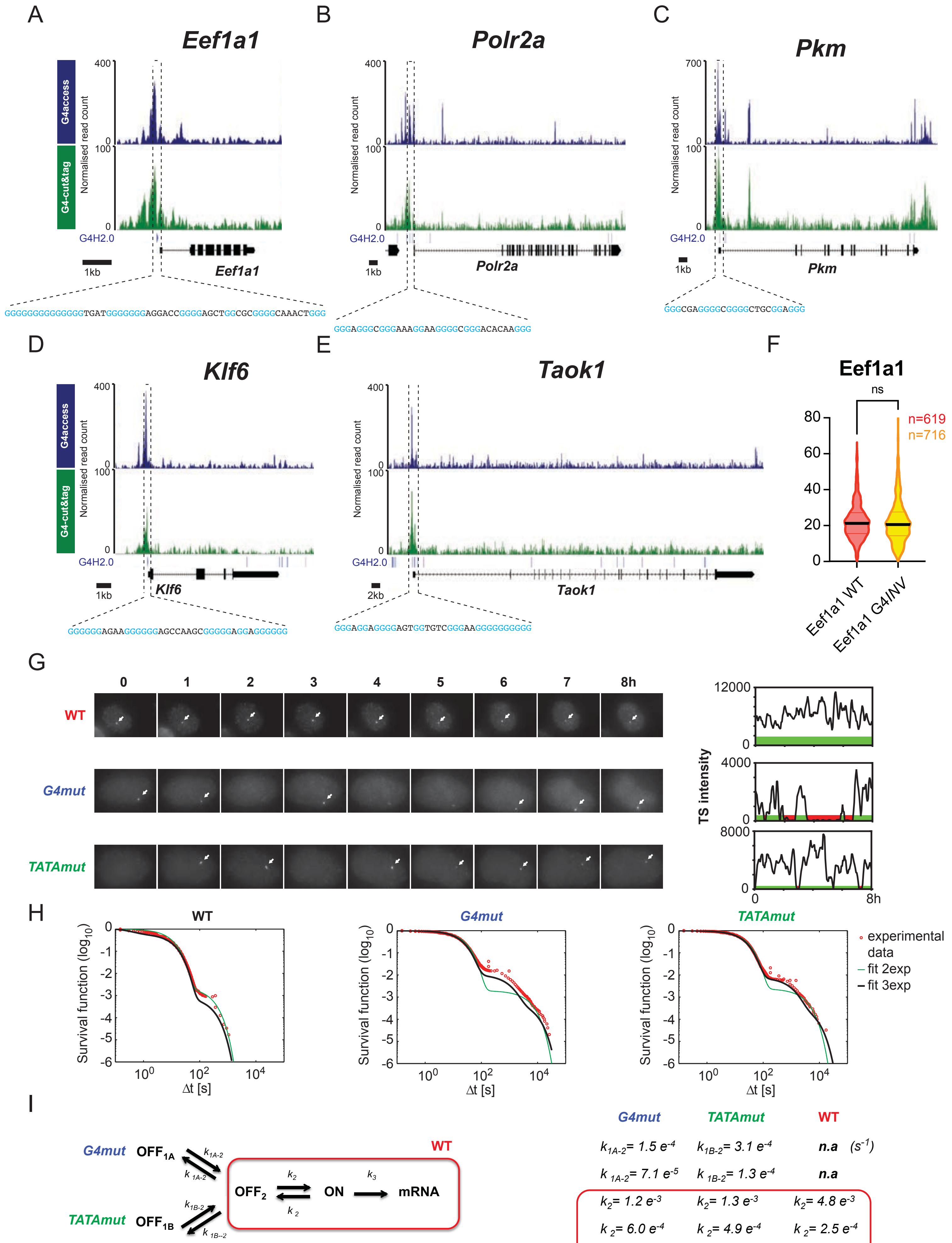


| G4mut | TATAmut | WT | |
|---|---|---|---|
| $k_{1A\text{-}2}$= 1.5 e$^{-4}$ | $k_{1B\text{-}2}$= 3.1 e$^{-4}$ | n.a | (s$^{-1}$) |
| $k_{\ 1A\text{-}2}$= 7.1 e$^{-5}$ | $k_{\ 1B\text{-}2}$= 1.3 e$^{-4}$ | n.a | |
| $k_2$= 1.2 e$^{-3}$ | $k_2$= 1.3 e$^{-3}$ | $k_2$= 4.8 e$^{-3}$ | |
| $k_{\ 2}$= 6.0 e$^{-4}$ | $k_{\ 2}$= 4.9 e$^{-4}$ | $k_{\ 2}$= 2.5 e$^{-4}$ | |
| $k_3$= 5.1 e$^{-2}$ | $k_3$= 5.4 e$^{-2}$ | $k_3$= 11.3 e$^{-2}$ | |

**Esnault *et al.*, Figure S7**

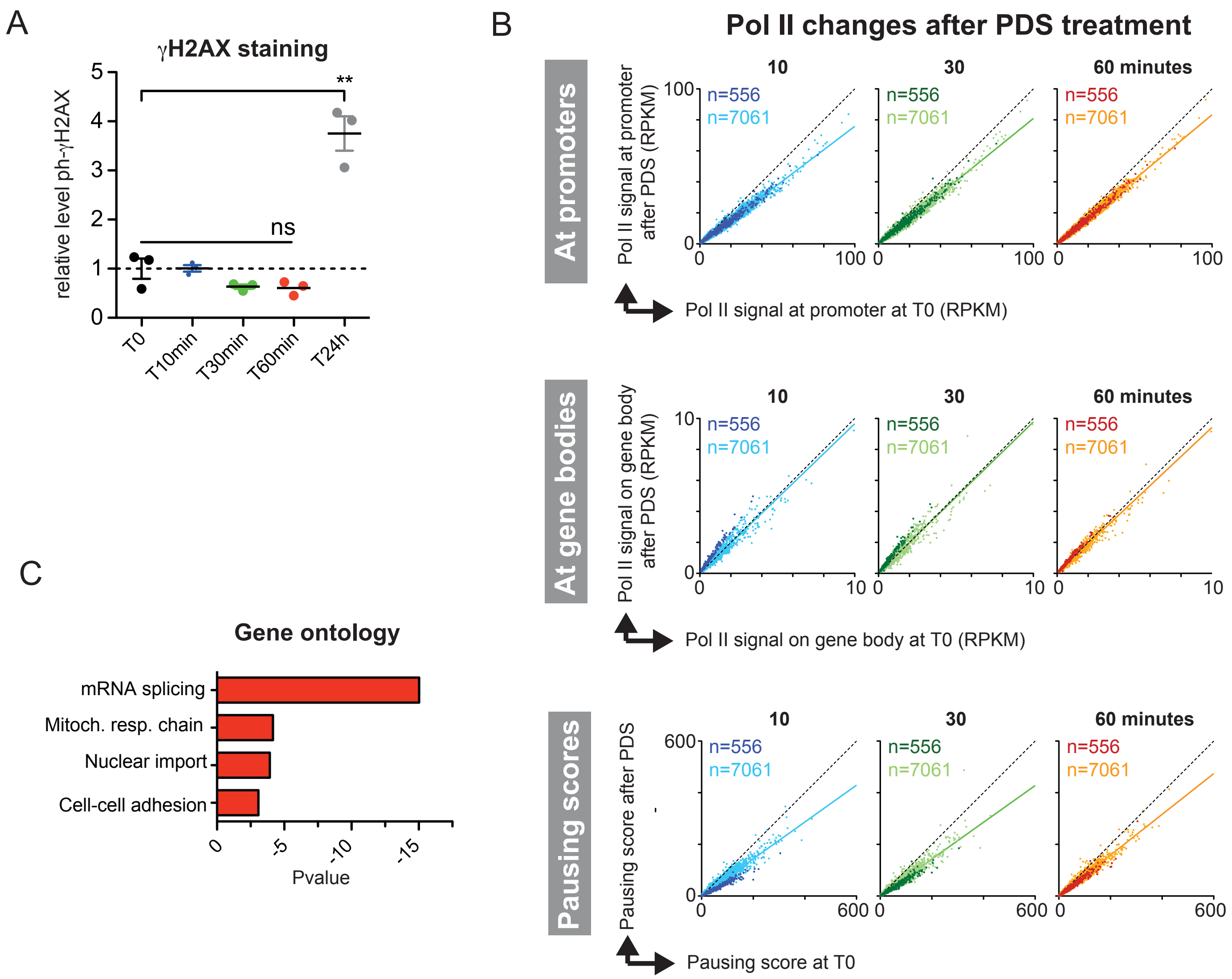
